# DNA conserved in diverse animals since the Precambrian controls genes for embryonic development

**DOI:** 10.1101/2023.06.18.545459

**Authors:** Martin C. Frith, Shengliang Ni

**Affiliations:** Artificial Intelligence Research Center, AIST; Graduate School of Frontier Sciences, University of Tokyo; Computational Bio Big Data Open Innovation Laboratory, AIST

## Abstract

DNA that controls gene expression (e.g. enhancers, promoters) has seemed almost never to be conserved between distantly-related animals, like vertebrates and arthropods. This is mysterious, because development of such animals is partly organized by homologous genes with similar complex expression patterns, termed “deep homology”.

Here we report twenty-five regulatory DNA segments conserved across bilaterian animals, of which seven are also conserved in cnidaria (coral and sea anemone). They control developmental genes (e.g. *Nr2f, Ptch, Rfx1/3, Sall, Smad6, Sp5, Tbx2/3*), including six homeobox genes: *Gsx, Hmx, Meis, Msx, Six1/2*, and *Zfhx3/4*. The segments contain perfectly or near-perfectly conserved CCAAT boxes, E-boxes, and other sequences recognized by regulatory proteins. More such DNA conservation will surely be found soon, as more genomes are published and sequence comparison is optimized. This reveals a control system for animal development conserved since the Precambrian.

## Introduction

Genes often remain similar across vast gulfs of evolution. For example, the genes that encode ribosomal RNA (rRNA) in humans and bacteria have recognizably-similar DNA sequences. Gene expression is controlled by DNA segments near the genes, which curiously lack such long-term conservation. Some are conserved across vertebrate animals, but few have been found conserved between vertebrates and invertebrates [1–3].

Early animal evolution produced bilaterian animals, which then split into two superphyla: protostomes and deuterostomes (fig. 1). A pioneering study by Royo et al. found several regulatory DNA segments conserved across deuterostomes, but found none of them in any non-deuterostome, except, amazingly, two in sea anemone, which is not even bilaterian [4]. These two are enhancers of the *Sox21* and *Hmx* genes. Soon after, Clarke et al. found two regulatory elements conserved between deuterostomes and protostomes [5]: an enhancer of the *Id* gene in gastropods (but no other protostomes), and an enhancer of *Znf503* in tick (but no other protostomes). Beyond this handful of exceptions, it seems that gene-regulating DNA is not conserved across bilaterians, suggesting that their developmental programs are not conserved either.

**Figure 1:**
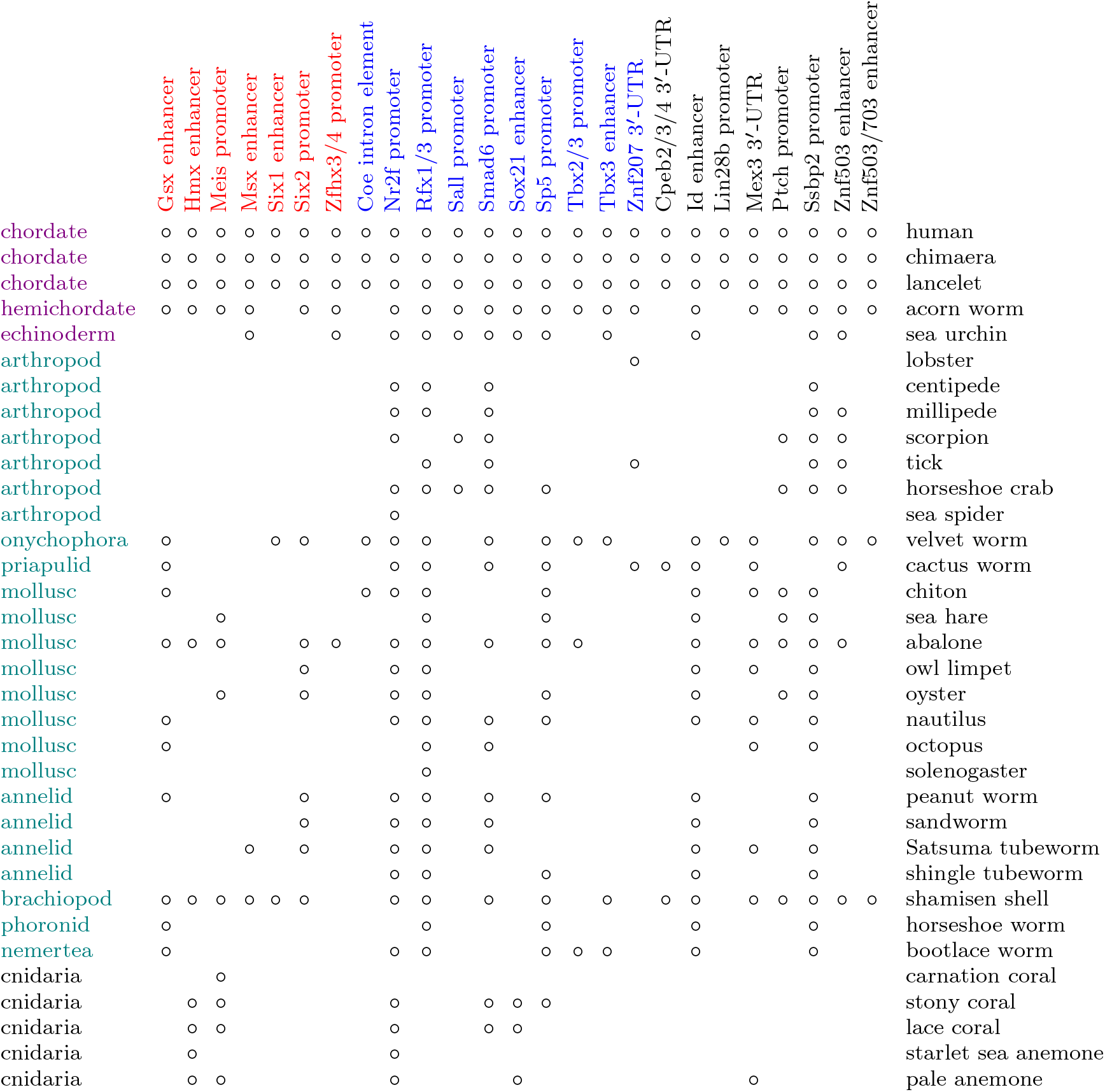
Animals in which each regulatory DNA element was detected. Regulatory segments are colored according to their nearby gene: homeobox genes red, other transcription factors blue. Each animal’s phylum is written on the left, deuterostomes in purple and protostomes in green.

Regulatory DNA conserved across vertebrates or beyond tends to regulate genes that control embryonic development [2]. These genes often encode transcription factors: proteins that control other genes by interacting with their regulatory DNA. One family of transcription factor genes contain homeobox sequences, which encode a DNA-binding structure called the homeodomain: these are especially used in development. Development involves the appearance of different territories in the embryo that express different combinations of transcription factors, and then develop into different body parts. These territories form in response to varying concentrations of signalling molecules, such as BMP, Wnt, and hedgehog proteins, that are secreted from cells. Often, one gene contributes to many parts of development, presumably because development evolved by redeploying these genes.

It is curious that developmental genes are widely conserved across animals, but their regulatory DNA rarely is. One theory is that conserved regulatory segments correspond to conserved body plans such as vertebrate [1]. This does not explain, however, why the expression patterns of developmental genes are often deeply conserved across animals [2]. One possible answer is that regulatory function can be conserved even as the DNA evolves so much that sequence similarity is lost [6].

Here, we find many more DNA segments that control developmental gene expression and are conserved across bilaterian animals, and even in non-bilaterian corals and sea anemone. They were found by exploiting the wealth of recently-published animal genomes, and by optimizing DNA homology detection [7]. This reveals a system of DNA sequences conserved since the Precambrian that control animal development.

## Results

The first step was to find conserved DNA segments between the genomes of human and chimaera, a cartilaginous fish. These genomes are related closely enough that many conserved segments can be found, but distantly enough that most of their DNA lacks similarity. Another reason for using chimaera is that its genome has evolved slowly [8]. Next, conserved regions were removed if they encode protein, rRNA, tRNA, snRNA, snoRNA, or miRNA. Pseudogenes were also removed, i.e. DNA that used to encode protein, rRNA, tRNA, etc. (Such DNA might also regulate gene expression, but its conservation can be explained by what it does or did encode.) The remaining conserved regions cover 5.7 million base-pairs in chimaera: 1/174-th of the genome.

The next step was to seek conserved segments between these chimaera regions and various invertebrate genomes. By not using the whole chimaera genome, we get 174-fold fewer false matches, so we can find 174-fold weaker similarities. Each similarity was given an *E*-value, which means the number of times such a similarity would be expected between random sequences of shuffled bases. More specifically, between random sequences with 40% g+c and the same lengths as: the one invertebrate genome, and all the chimaera regions. Similarities were accepted with either *E*-value ≤0.0001, or *E*-value ≤10 and near homologous genes. For example, a chimaera segment near the *smad6* gene matched octopus DNA with *E*-value 0.46, and the nearest gene in octopus is the homolog of *smad6*.

As a negative control, each search was repeated after reversing (but not complementing) the invertebrate genome. DNA doesn’t evolve by reversal, because that would require flipping the 3^*I*^-to-5^*I*^ bonds between all the nucleotides. In most cases a reversed genome had about 10 matches with *E*-value ≤ 10, and the lowest *E*-value ever seen was ≤ 0.0009.

Further sensitivity was gained by using conserved parts of invertebrate genomes. For example, by using oyster DNA segments that are conserved in scallop (1/59-th of the oyster genome), sensitivity was boosted a further 59-fold, revealing matches near the *Ptch* and *Meis* genes.

These methods found 25 DNA segments with similarity between vertebrates and non-deuterostome animals (fig. 1). They can be explained by either common ancestry or convergent evolution. It seems implausible that most are due to convergent evolution, because they are in a wide range of animals (fig. 1, 2) and they preserve position and orientation relative to the genes (fig. 3). The segments were not found in some invertebrates (fig. 1): either they were lost during evolution of those animals, or they are present but undetected. The segments are labelled “promoter” if they occur at a gene’s transcription start site in human (the best-annotated genome), else “enhancer”. One segment lies in an intron, and three lie in 3^*I*^-UTRs (UnTranslated Regions of exons, downstream of the translated protein-coding region).

**Figure 2:**
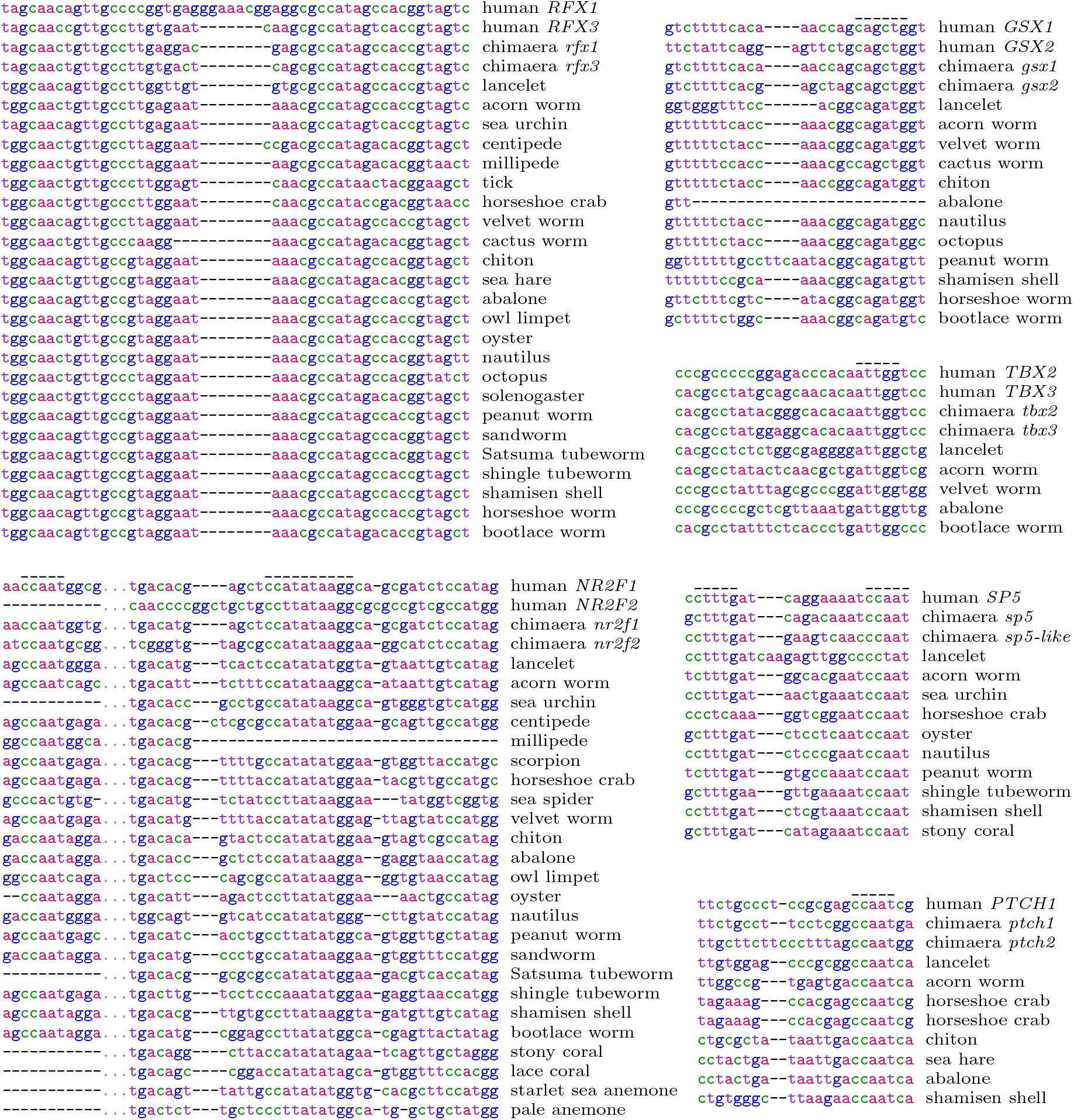
Parts of alignments between regulatory DNA sequences. The full alignments are in the Supplement. The over-bars show: CCAAT boxes (in the *Nr2f, Sp5*, and *Ptch* promoters), reverse-strand CCAAT box ATTGG (*Tbx2/3* promoter), E-box CAnnTG (*Gsx* enhancer), Wnt response element CTTTG (*Sp5* promoter), and CArG box CC(A/T)_6_GG (*Nr2f* promoter). The *Ptch* promoter matches two places in the horseshoe crab genome, both near genes annotated as “patched-like”.

**Figure 3:**
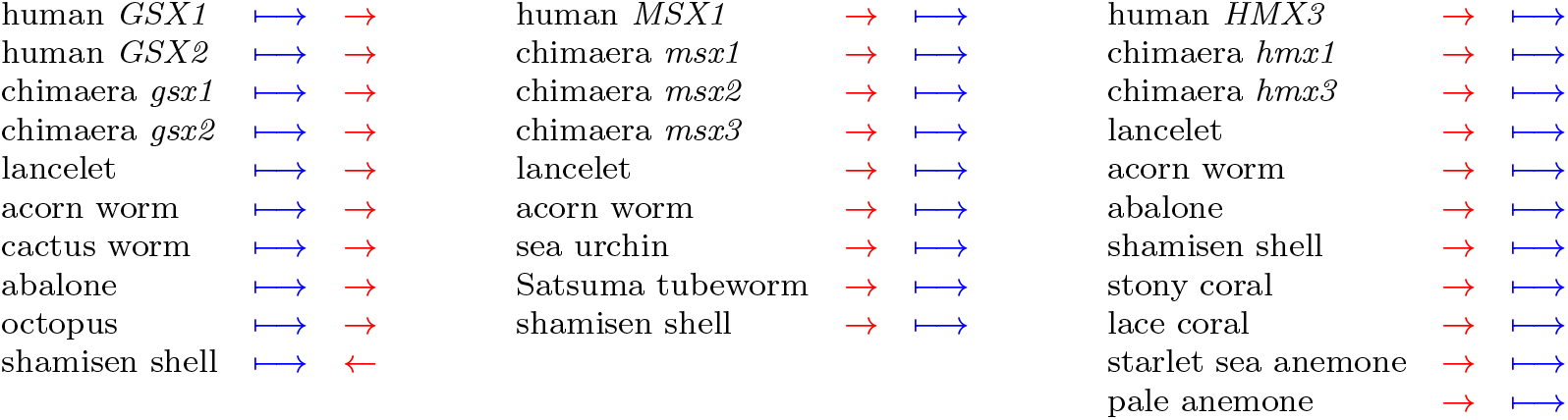
Position and orientation of conserved DNA segments (red arrows) relative to nearby genes (blue arrows). Within each column, all the genes are homologous to each other. Animals without gene annotations are not shown. A regulatory segment doesn’t really have an orientation, but once an orientation is arbitrarily chosen for any one segment, that defines the orientations of all its homologs.

Interestingly, all of these genes, except *Znf207*, have multiple (2–4) copies in genomes of jawed vertebrates. For example, jawed vertebrates have three similar *Meis* genes (*Meis1, Meis2*, and *Meis3*), two *SoxB2* genes (called *Sox14* and *Sox21*), two *sine oculis* genes (*Six1* and *Six2*), and two *NET* genes (*Znf503* and *Znf703*). *Sp5* is a semi-exception: vertebrates have two copies, named *Sp5* and *Sp5-like*, but *Sp5-like* was lost in mammals and birds [9]. In early vertebrate evolution, there were two rounds of whole-genome duplication. Most gene duplicates were rapidly lost, so that vertebrates do not have many more genes than invertebrates, but some genes remain in 2–4 copies, including nearly all the ones here.

The conserved DNA segments were detected at all, some, or just one of these multi-copy genes. For example, the conserved *Gsx* enhancer was found near both *Gsx* genes in human and chimaera (fig. 2, 3), but the *SoxB2* enhancer was only found at *Sox21*. The *sine oculis* enhancer was only found at *Six1*, whereas the *sine oculis* promoter was only found at *Six2*. One *NET* enhancer was found at both *Znf503* and *Znf703*, the other only at *Znf503*. While “not found” does not necessarily mean “absent”, it seems that duplication of these genes enabled divergence of their regulation and developmental roles.

The functions of these genes and conserved DNA segments are partially known. *Gsx* and *Msx* encode transcription factors that define territories in early development of the CNS (central nervous system), in a manner conserved across bilaterian animals [10–12]. They produce an ancient sensory/inter/motor neuron system. *Msx* is expressed at the lateral edges of the CNS, and produces Rohon–Beard mechanosensory neurons. *Gsx* is expressed more medially and produces interneurons, and the most medial territory produces motor neurons. The *Gsx* enhancer seems to contribute by preventing lateral expression of *Gsx*, thus defining the brain’s pallium-subpallium boundary [13, 14]. *Msx* plays many other roles in development: the enhancer is the previously-identified *Msx1* “proximal element” that produces expression in mouse embryo nasal epithelium, myotome, limb mesenchyme, eye, ear, roof plate, second arch, genital ridge and epiphysis [15].

The *Six1/2* genes control development of eyes and other cranial sensory organs (e.g. olfactory epithelium, inner ear, taste papillae), and also contribute to development of kidney, thymus, parathyroid gland, and more [16]. The *Six1* enhancer is the previously-described Six1-13 element, which produced variable expression patterns of unclear importance [16]: it would be interesting to re-examine this element.

*Tbx2/3* control formation of the brain’s neurohypophysis [17], and contribute to development of vertebrate limbs, heart, liver, and more [18]. The *Tbx3* enhancer was shown to affect expression of *Tbx3* and limb length in mice and horses [19].

The *Coe* genes encode transcription factors affecting subpallium development, expressed in *Gsx* + brain regions of mice and annelids [10]. They also define subtypes of body-wall motor neuron in mice and roundworms [20], and promote development of fat cells [21]. The conserved DNA segment is in an intron, so is transcribed into RNA: it might function in the DNA or RNA or both. This RNA region, of the mouse *Coe* gene *EBF1*, interacts with PSPC1 protein during fat cell development, promoting export of the RNA from the nucleus and expression of EBF1 protein [21].

The *Znf503/703* genes encode transcriptional repressors that are related to Sp transcription factors, but are thought not to bind directly to DNA. They contribute to brain and eye development in vertebrates and flies. In vertebrates they define hindbrain territories and motor neuron subtypes, and contribute to development of the brain’s striatum, and limbs [22]. The *Znf503* enhancer was previously found in tick [5]: here it is found in diverse protostomes (fig. 1). The *Znf503/703* enhancer is 2.5 kb further upstream: it directs expression, in mouse embryos, in branchial arches and their derivatives, apical ectodermal ridge of limb buds, and urogenital tissues [23].

The other genes also regulate development [24–27]. *Hmx* interacts with *Gsx* in development of the brain’s hypothalamus [28]. *Ptch* genes encode the main receptors that recognize hedgehog signalling molecules, and Smad6 proteins inhibit response to BMP signalling molecules [29]. *Sp5* is a target of Wnt signalling in bilaterians, and also in cnidaria where it prevents development of multiple heads [30]. It has a promoter region conserved from human to coral, which includes an especially conserved sequence resembling a Wnt response element (fig. 2). The *Rfx1/3* genes control formation of cilia on cells, which are used for hedgehog signalling, establish left-right body asymmetry, and more [31]. The promoter is spectacularly conserved across bilateria (fig. 2). Ssbp2 protein partners with LIM homeodomain proteins, which specify motor neurons in the same way in vertebrates and flies, among other roles [32]. *Nr2f* participates in neural development of bilaterians and cnidaria [33]: its promoter has been conserved since early neural evolution (fig. 1, 2).

Interestingly, the three genes with conserved 3^*I*^-UTR segments encode proteins that bind to RNA. The Cpeb proteins control mRNA localization and translation, in neural development and memory formation [34]. They are regulated by miRNAs binding to their 3^*I*^-UTRs [35–37]. *Mex3* contributes to defining anterior territories in bilaterian embryos, by destabilizing mRNAs [38, 39]. Its 3^*I*^-UTR contains elements for translational enhancement and destabilization, including by its own protein or miRNA [38, 40]. The UTR was previously found conserved between vertebrates and lancelet [38]: here it is found in other phyla and even a sea anemone (fig. 1). The Znf207 protein is a bit of an outlier: it contributes to chromosome alignment during cell division, a function not specific to animal development. It also, however, contributes to early neural development by binding to DNA or RNA [41].

Do these particular developmental genes have a common theme? It is hard to say, but perhaps there is a tendency for roles in ventral CNS (e.g. subpallium, striatum, hypothalamus, neurohypophysis, motor neurons).

In the conserved DNA segments, conserved transcription factor binding sites were identified (see the Supplement), using the JASPAR database of DNA-binding tendencies [42]. The results are uncertain, because transcription factors have weak sequence preferences, often similar to those of other transcription factors. Perhaps different factors bind to overlapping sites at different times depending on molecule concentrations.

On the other hand, there are clearly-conserved “boxes”, including E-boxes and many CCAAT boxes (fig. 2). The E-box, CAnnTG, binds to various transcription factors that have a bHLH (basic-helix-loop-helix) structure. bHLH factors initiate neural development in bilateria and cnidaria [11], perhaps by binding to these conserved E-boxes. CCAAT boxes bind to the transcription factor NF-Y, which controls many developmental genes [43]. One CArG box was found, conserved from human to sea anemone (though imperfect in coral), in the *Nr2f* promoter (fig. 2). In general, CArG boxes bind to the Srf transcription factor, which controls cell migration and neural outgrowth in development [44].

Another interesting result is that the conserved DNA segments rarely change position (upstream/downstream) or orientation relative to the nearby genes. Figure 3 shows just one change, for *Gsx* in shamisen shell. There are a few others. In chimaera, the *sox21* gene has been inverted relative to its neighboring genes, but the inverted region doesn’t include the enhancer. So, the enhancer is upstream of *sox21* in chimaera but downstream in the other animals.

Some protostome animals have more detected DNA elements than others (fig. 1). This is partly artifactual, e.g. some genomes lacked gene annotations so had the stricter 0.0001 *E*-value threshold. With that said, these seem to have many conserved elements: abalone, shamisen shell, and velvet worm (which lacks gene annotations). Perhaps these animals have undergone less evolution of their development and body plan. All three have sometimes been regarded as “primitive” or “living fossils”: the shamisen shell (*Lingula*) was noted in Darwin’s Origin of Species as little changed from ancient fossils.

On the other hand, no conserved segments were found in fly (*Drosophila melanogaster*), round-worm (*Caenorhabditis elegans*), or the leech *Helobdella robusta*. Perhaps the developmental systems of these animals have evolved more drastically [45]. Beyond bilateria, no conserved segments were found in ctenophores (comb jellies), sponges, or *Trichoplax* (Supplement table 2). A recent study found three regulatory DNA elements conserved between mammals and flies [46]: we confirmed they are conserved between human and chimaera, but didn’t find them in invertebrates. Even when comparing each pair of human and fly sequences, boosting sensitivity 10^12^-fold over comparing the whole genomes, our methods found tiny similar fragments in just one out of three pairs. Another study found identical DNA segments ≥ 50 bp between human and fly, and even human and sponge [47]. We confirmed that such intriguing matches exist, though they overlap regions that encode protein, rRNA, tRNA, or snRNA, or pseudogenes. A third study found 6 533 non-protein-coding segments conserved across deuterostomes [48], vastly more than found here or elsewhere [4, 5]. Some possible reasons are that they used *E*-value ≤ 0.05 per 5 kb (which is 1 per 100 kb), and didn’t exclude pseudogenes or RNA genes.

In summary, we begin to see a system of DNA elements controlling gene expression for animal development, conserved since ancestral bilaterians and beyond in the Precambrian. This conservation helps us understand how early animals developed, and how modern development is built on these ancient components.

Why have these DNA sequences been so conserved? Judging by present-day bilaterians and cnidaria, the common ancestor of these animals already had an exquisitely integrated multicellular body, with a nervous system, that developed from a single fertilized cell [11, 12]. This complex development must have evolved by accretion of developmental processes, often by redeploying existing pathways [28]. So, one regulatory DNA element could become used in many parts of development, or its outcomes might get built upon by many processes. This brings us to Hyrum’s Law of software development: “With a sufficient number of users… all observable behaviors of your system will be depended on by somebody” [49]. Thus, DNA changes with even subtle effects on transcription factor interactions, e.g. though DNA shape [50], would likely be harmful. This resulted in strongly-conserved DNA controlling core animal development, upon which evolved arthropods, molluscs, chordates, and so on.

