## Supplement for "DNA conserved in diverse animals since the Precambrian controls genes for embryonic development"

Martin C. Frith[1,2,3] and Shengliang Ni[2]

[1]Artificial Intelligence Research Center, AIST
[2]Graduate School of Frontier Sciences, University of Tokyo
[3]Computational Bio Big Data Open Innovation Laboratory, AIST

#### Data availability

The alignments between pairs of whole genomes (e.g. chimaera-human, oyster-scallop) are in the 2023 section of `https://github.com/mcfrith/last-genome-alignments`. Other files used in this study are at `https://github.com/mcfrith/conserved-animal-dna`.

#### Methods

##### Finding conserved segments between the human and chimaera genomes

The genomes were compared using LAST version 1453 (`https://gitlab.com/mcfrith/last`). First, `chrEBV` (Epstein-Barr virus) was removed from the human genome. An index called (say) `cdb` was made for the chimaera genome:

```
lastdb -P8 -uMAM8 -c cdb chimaera.fa
```

`-P8` makes it faster by using 8 processors, `-uMAM8` increases sensitivity and memory use [1], and `-c` suppresses simple sequences like `atatatatatatatatat`. Next, rates of insertion, deletion, and substitutions between the genomes were found [2]:

```
last-train -P8 --revsym -D1e9 --sample-number=5000 cdb human.fa > hc.train
```

Then, many-to-one human-to-chimaera alignments were found (based on these rates) [3]:

```
lastal -P8 -D1e9 -m100 --split-f=MAF+ -p hc.train cdb human.fa > many-to-one.maf
```

`-m100` makes it more slow and sensitive, and `-D1e9` gets similarities that would be expected between shuffled-base sequences at most once per $10^9$ human bp. The alignments were cut down to one-to-one alignments [3]:

```
last-split -r many-to-one.maf > one-to-one.maf
```

Finally, isolated alignments (not near other alignments) were discarded, using `maf-linked` from `https://gitlab.com/mcfrith/protein-fossils/`:

```
maf-linked one-to-one.maf > linked.maf
```

This final step removes non-homologous insertions of homologous transposons [4].

#### Human genes and pseudogenes

In order to find conserved regulatory DNA, we need to exclude conserved DNA in protein-coding regions, RNA genes, and pseudogenes. The following genome coordinates were used, where `.txt` and `.bigPsl` refer to files in the UCSC genome database:

- Protein-coding regions in `ncbiRefSeq.txt`, `wgEncodeGencodeCompV43.txt`, `xenoRefGene.txt`, and `hg38.refseq.transMapV5.bigPsl`.
- rRNA, tRNA, and snRNA genes and pseudogenes in `rmskOutCurrent.txt`.
- Exons of RNA genes in `ncbiRefSeq.txt`, extracted like this:

  ```
  awk '$13 ~ /^RNU|^MIR|^SNOR|^SCARNA|^RNA5-8/' ncbiRefSeq.txt
  ```

- Protein fossils derived from host genes, in `https://github.com/mcfrith/protein-fossils`.

#### Chimaera genes and pseudogenes

First, repeats were found with RepeatMasker 4.1.5 (using RMBlast 2.13.0 and TRF 4.09):

```
RepeatMasker -xsmall -engine rmblast -s -species "Callorhinchus milii" chimaera.fa
```

Then, protein fossils were found as described previously [4], except that `fasta-nr` option `-s` was used this time. Finally, these genome coordinates were obtained:

- Protein-coding regions and RNA genes in the genome's `genomic.gff` file, extracted like this:

  ```
  awk -F"\t" '$3 ~ /CDS|rRNA|tRNA|snRNA|snoRNA|guide_RNA/' genomic.gff
  ```

  Here, `guide_RNA` means scaRNA (small Cajal body-specific RNA), which is a type of snoRNA.
- rRNA, tRNA, and snRNA genes and pseudogenes from RepeatMasker.
- Protein fossils derived from host genes.

#### Non-genic chimaera DNA conserved in human

First, each human (pseudo)gene segment was extended by 50 base-pairs in both directions. This is because the edges of the human-to-chimaera alignments cannot be perfectly reliable: they may overshoot slightly beyond conserved regions into non-conserved DNA, or undershoot [5]. Then, chimaera bases were retained if they align to human bases not in these segments. These chimaera regions were extended by 50 bases in both directions. Finally, bases in chimaera (pseudo)genes were removed from these regions.

#### Comparing invertebrate genomes to these chimaera regions

An index called (say) `sdb` was made for the chimaera DNA regions:

```
lastdb -c -uMAM8 -S2 sdb chimaera-segments.fa
```

Then, similar segments were sought between each genome and the chimaera regions:

```
lastal -P8 -J1 -D$D -m100 -p my.train sdb genome.fa > out.maf
```

Here, `$D` was set to (genome size) / 10: this gets similarities that would be expected at most once per `$D` genome base-pairs if we compared a shuffled genome to shuffled DNA with the same length as the whole of `sdb`. In other words, it gets similarities with $E$-value $\leq 10$. This step requires a file `my.train`, which is explained later.

#### $E$-value adjustment

For each pair of similar segments, `lastal` outputs a per-sequence $E$-value. This is the number of times such a similarity would be expected between random DNA sequences with the same lengths as: the one sequence from `genome.fa` (chromosome or fragment), and the whole of `sdb`. These $E$-values were converted to per-genome $E$-values:

genome $E$-value = (sequence $E$-value) × (genome length) / (sequence length) .

#### Simple sequences and reversed genomes

Simple sequences, like `atatatatatatatatat`, evolve frequently and independently. This leads to similar segments that are not conserved homologs. There are several methods to suppress simple sequences from homology search, but these methods do not work equally well. LAST uses the `tantan` method, which seems to suppress them effectively [6].

One way to test this is by comparing two DNA sequences after reversing (but not complementing) one of them. Then there are no true homologs, but simple sequences produce many strong similarities [6]. In the present study, reversed genomes did not have much stronger matches than would be expected between random sequences of shuffled bases, indicating that simple sequences were suppressed effectively.

#### Finding conserved regions between invertebrate genomes

This step found, for example, oyster DNA segments that are conserved in scallop. The genomes were compared in the same way as human-to-chimaera, without the final `maf-linked` step. Oyster protein-coding regions and RNA genes were extracted like this:

```
awk -F"\t" '$3 ~ /CDS|rRNA|tRNA|snRNA|snoRNA|guide_RNA/' genomic.gff
```

Each of these regions was extended by 50 base-pairs in both directions. Then, these regions were cut from the oyster segments that align to scallop. The remaining oyster segments were then extended by 50 bases in both directions.

This was repeated for the genome pairs in table 4, with oyster replaced by the left genome and scallop by the right genome. Some of these genomes did not have a `genomic.gff` file, so an empty file was used instead.

#### The `my.train` file

This file has rates of insertion, deletion and substitutions. It defines "similar": for example, it might specify that `a`↔`g` and `c`↔`t` substitutions are more likely than other substitutions.

Typically, `last-train` finds these rates from the sequences we wish to compare [2]. But that doesn't work here, because the invertebrate DNA is too distantly related to the chimaera regions. Therefore, a more closely-related animal, lamprey (a jawless vertebrate), was used instead. Using the whole lamprey genome doesn't work well either, so lamprey DNA segments were used that are conserved in hagfish (another jawless vertebrate). These conserved lamprey segments were found in the same way as oyster-versus-scallop, removing protein-coding regions and RNA genes in the same way too. These lamprey segments were compared to chimaera in two ways:

```
last-train -s1 --revsym --matsym --gapsym -D1e7 --sample-number=5000
                                              sdb lampreySegs.fa > try1.train

last-train -s1 --revsym --matsym --gapsym -D1e7 --sample-number=5000 --pid=65 --scale=6
                                              sdb lampreySegs.fa > try2.train
```

`--pid=65` makes `last-train` ignore similarities with > 65% identity, so the rates will be tuned for finding homologies with low percent-identity, but will be worse for finding short homologies with high percent-identity. (`--scale=6` may be unnecessary: it makes the output have units of 1/6 bits). Both of these `train` files were used for homology searches.

#### Finding matches to conserved deuterostome DNA

After comparing invertebrate DNA to the chimaera regions, there were 45 chunks of the chimaera regions conserved in at least one invertebrate. Most of these chunks are conserved in lancelet or acorn worm.

To boost sensitivity further, the searches were repeated against these chunks of chimaera, lancelet, and acorn worm. The chunks were first extended by 50 bases in both directions. Then, protein-coding regions and RNA genes of the three animals were extracted like this:

```
awk -F"\t" '$3 ~ /CDS|rRNA|tRNA|snRNA|snoRNA|guide_RNA/' genomic.gff
```

and cut from the conserved chunks. The chunks were indexed:

```
lastdb -c -uMAM8 -S2 ddb chimaera-lancelet-acornWorm-chunks.fa
```

Then similarities were found using `lastal` as above.

Thus, the searches were against deuterostome DNA (from chimaera, lancelet, and acorn worm). In future, more DNA conservation can probably be found by using non-deuterostomes too. The present study stopped here, to avoid risk of "transitive catastrophe": see for example stony coral and bootlace worm in the *Sp5* promoter alignment.

#### Multiple sequence alignment

In the end, after finding the conserved segments, the DNA sequences were aligned with the `linsi` method in MAFFT v7.520 [7]:

```
linsi sequences.fa > alignment.fa
```

#### Transcription factor binding signals

The DNA sequences were scanned with transcription factor binding site profiles from the JASPAR database (non-redundant vertebrate profiles in JASPAR CORE 2022) [8]. The scanning was done by FIMO version 5.5.2 [9].

#### miRNA target signals

Potential miRNA binding sites were found with TargetScan Release 8.0 [10].

### Tables

**Table 1:** Genomes with detected regulatory elements (size means number of a/c/g/t bases in the file)

| Animal | Phylum | Scientific name | Genome | Size (bp) | Has gene annotation |
|---|---|---|---|---|---|
| human | chordate | *Homo sapiens* | `hg38_no_alt_analysis_set` | 2934876451 | yes |
| chimaera | chordate | *Callorhinchus milii* | `GCF_018977255.1` | 991467956 | yes |
| lancelet | chordate | *Branchiostoma floridae* | `GCF_000003815.2` | 487128766 | yes |
| acorn worm | hemichordate | *Saccoglossus kowalevskii* | `GCF_000003605.2` | 642223636 | yes |
| sea urchin | echinoderm | *Strongylocentrotus purpuratus* | `GCF_000002235.5` | 921518293 | yes |
| lobster | arthropod | *Homarus americanus* | `GCF_018991925.1` | 2291242282 | yes |
| centipede | arthropod | *Strigamia maritima* | `GCA_000239455.1` | 173599458 | |
| millipede | arthropod | *Glomeris maerens* | `GCA_023279145.1` | 149766888 | |
| scorpion | arthropod | *Centruroides sculpturatus* | `GCF_000671375.1` | 863041242 | yes |
| tick | arthropod | *Ixodes scapularis* | `GCF_016920785.2` | 2226616818 | yes |
| horseshoe crab | arthropod | *Limulus polyphemus* | `GCF_000517525.1` | 1705786612 | yes |
| sea spider | arthropod | *Nymphon striatum* | `GCA_016618385.1` | 744500873 | yes |
| velvet worm | onychophora | *Euperipatoides rowelli* | `GCA_003024985.2` | 1744674727 | |
| cactus worm | priapulid | *Priapulus caudatus* | `GCF_000485595.1` | 436582939 | yes |
| chiton | mollusc | *Acanthopleura granulata* | `GCA_016165875.1` | 544712841 | |
| sea hare | mollusc | *Aplysia californica* | `GCF_000002075.1` | 737797481 | yes |
| abalone | mollusc | *Haliotis rufescens* | `GCF_023055435.1` | 1334438814 | yes |
| owl limpet | mollusc | *Lottia gigantea* | `GCF_000327385.1` | 298894987 | yes |
| oyster | mollusc | *Crassostrea gigas* | `GCF_902806645.1` | 647867327 | yes |
| nautilus | mollusc | *Nautilus pompilius* | `GCA_018389105.1` | 729019367 | |
| octopus | mollusc | *Octopus bimaculoides* | `GCF_001194135.2` | 1986279820 | yes |
| solenogaster | mollusc | *Wirenia argentea* | `GCA_025802215.1` | 528706996 | |
| peanut worm | annelid | *Sipunculus nudus* | `GCA_026874595.1` | 1426680931 | |
| sandworm | annelid | *Alitta virens* | `GCA_932294295.1` | 671131125 | |
| Satsuma tubeworm | annelid | *Lamellibrachia satsuma* | `GCA_022478865.1` | 664980950 | yes |
| shingle tubeworm | annelid | *Owenia fusiformis* | `GCA_903813345.2` | 499118861 | yes |
| shamisen shell | brachiopod | *Lingula anatina* | `GCF_001039355.2` | 389003419 | yes |
| horseshoe worm | phoronid | *Phoronis ovalis* | `GCA_028565635.1` | 325077420 | |
| bootlace worm | nemertea | *Lineus longissimus* | `GCA_910592395.2` | 391166316 | |
| carnation coral | cnidaria | *Dendronephthya gigantea* | `GCF_004324835.1` | 286150628 | yes |
| stony coral | cnidaria | *Acropora millepora* | `GCF_013753865.1` | 475342477 | yes |
| lace coral | cnidaria | *Pocillopora damicornis* | `GCF_003704095.1` | 225744106 | yes |
| starlet sea anemone | cnidaria | *Nematostella vectensis* | `GCF_932526225.1` | 269383238 | yes |
| pale anemone | cnidaria | *Exaiptasia diaphana* | `GCF_001417965.1` | 210157303 | yes |

**Table 2:** Genomes analyzed but no regulatory elements detected

| Animal | Phylum | Scientific name | Genome | Size (bp) | Has gene annotation |
|---|---|---|---|---|---|
| fruit fly | arthropod | *Drosophila melanogaster* | GCF_000001215.4 | 142573024 | yes |
| bee | arthropod | *Apis mellifera* | GCF_003254395.2 | 223937270 | yes |
| bristletail | arthropod | *Machilis hrabei* | GCA_003456935.1 | 1320889507 | |
| roundworm | nematode | *Caenorhabditis elegans* | GCF_000002985.6 | 100286401 | yes |
| jawless leech | annelid | *Helobdella robusta* | GCF_000326865.1 | 215435648 | yes |
| planarian | platyhelminth | *Schmidtea mediterranea* | GCA_022537955.1 | 773867583 | |
| flatworm | platyhelminth | *Macrostomum lignano* | GCA_002269645.1 | 762829307 | yes |
| brown bryozoan | bryozoa | *Bugula neritina* | GCA_010799875.2 | 214708255 | yes |
| hydra | cnidaria | *Hydra vulgaris* | GCF_022113875.1 | 817163865 | yes |
| trichoplax | placozoa | *Trichoplax adhaerens* | GCF_000150275.1 | 94748975 | yes |
| sea walnut | ctenophore | *Mnemiopsis leidyi* | GCA_000226015.1 | 150338246 | |
| comb jelly | ctenophore | *Bolinopsis microptera* | GCA_026151205.1 | 265424760 | |
| sea gooseberry | ctenophore | *Hormiphora californensis* | GCA_020137815.1 | 110660632 | |
| cigar comb jelly | ctenophore | *Beroe ovata* | GCA_946803715.1 | 84911038 | |
| demosponge | porifera | *Amphimedon queenslandica* | GCF_000090795.2 | 143136305 | yes |
| glass sponge | porifera | *Oopsacas minuta* | GCA_024704765.1 | 60959430 | yes |

**Table 3:** Other genomes used in this study

| Animal | Phylum | Scientific name | Genome | Size (bp) | Has gene annotation |
|---|---|---|---|---|---|
| lamprey | chordate | *Petromyzon marinus* | GCF_010993605.1 | 1074036726 | yes |
| hagfish | chordate | *Eptatretus burgeri* | GCA_024346535.1 | 1507324335 | |
| Bahama lancelet | chordate | *Asymmetron lucayanum* | GCA_001663935.1 | 452764958 | |
| Hawaiian acorn worm | hemichordate | *Ptychodera flava* | GCA_001465055.1 | 1126321642 | |
| sea cucumber | echinoderm | *Apostichopus japonicus* | GCA_002754855.1 | 798993713 | yes |
| robber fly | arthropod | *Machimus atricapillus* | GCA_933228815.1 | 268640068 | |
| woodlouse | arthropod | *Armadillidium nasatum* | GCA_009176605.1 | 1223004717 | yes |
| sea butterfly | mollusc | *Limacina bulimoides* | GCA_009866985.1 | 2900980226 | |
| top snail | mollusc | *Phorcus lineatus* | GCA_921293015.1 | 957783961 | |
| scallop | mollusc | *Mizuhopecten yessoensis* | GCF_002113885.1 | 907590157 | yes |
| firefly squid | mollusc | *Watasenia scintillans* | GCA_015471945.1 | 649114004 | |
| medicinal leech | annelid | *Hirudo medicinalis* | GCA_011800805.1 | 156118814 | |
| waratah anemone | cnidaria | *Actinia tenebrosa* | GCF_009602425.1 | 214400470 | yes |
| zoanthid | cnidaria | *Epizoanthus planus* | GCA_025388665.1 | 215910112 | |

**Table 4:** Invertebrate genomes that were compared to each other

| Given to `lastdb` | Given to `lastal` |
|---|---|
| stony coral | waratah anemone |
| sea hare | sea butterfly |
| lancelet | Bahama lancelet |
| lancelet | acorn worm |
| oyster | scallop |
| fruit fly | robber fly |
| millipede | centipede |
| abalone | top snail |
| jawless leech | medicinal leech |
| lobster | woodlouse |
| Satsuma tubeworm | shingle tubeworm |
| horseshoe crab | scorpion |
| shamisen shell | horseshoe worm |
| sea walnut | sea gooseberry |
| starlet sea anemone | zoanthid |
| octopus | firefly squid |
| acorn worm | Hawaiian acorn worm |
| sea urchin | sea cucumber |

### Alignments

Above each alignment are shown transcription factor binding motifs that are conserved in most of the sequences. Some are labeled by name of the (family of) transcription factor. Others are labeled by "box" or "element":

- E-box: `canntg`, bound by many bHLH (basic helix-loop-helix) transcription factors and some zinc finger transcription factors.
- CCAAT: CCAAT-box, bound by NF-Y.
- WRE: Wnt response element, bound by TCF/LEF proteins, the consensus is said to be `ctttg` or `cctttgww`.
- NRRE: nuclear receptor response element, the consensus is said to be `aggtca` or `(a/g)g(g/t)tca`.
- GC-box: bound by Sp transcription factors, the consensus is `gggcggg` or similar.
- CArG-box: `ccwwwwwwgg`, bound by transcription factor Srf.

```
                                                                                                                                                          Sox___
human GSX1          --gggtctgtta-------gacggtgcggtgcctgcagctgatttgatgggttggaaatgcgcacgaaacaaaatacttgaat-cctgaaaaagaccctttgtaagtttctt-tagaaatcaagcaattctgcatgacaa
human GSX2          --gggtctggca----ggcgcagaggtgccgtatgcaaatggttc------actattaggcgagaaaga-----------aatacttcaaatggccctttgtaaccttctat-agaaaatcgaggtactctgcatgacaa
chimaera gsx1       -----tctggta-------gacggtgcgatgcacgcagctgatttgatgagttggaaatacgcacgaa--aaaatacttgaat-cctgaaaaagatcctttgtaagtttctt-tagaaatcaagcaattctgcatgacaa
chimaera gsx2       --gggtctggta-------ggcgaggtgggatatgcaaatga-----------caggatattcgcaaggaaaaatactcgaatcctccaaatggctcttttgtgagtttctt-tagaaatcaagcaattctgcatgacaa
lancelet            cccaacctgctgtcggggggcagagataattttccaaaatggagt-----ggaaaatagacc--------aaaatatatagca-gcgcgccgggctttgtgctgcttgtatataagaaaacaacatattctacatgacaa
acorn worm          cgcaatctgctg--gccgcagccagataatttatgcaaatga-----------ggcgcggcaacgccag-aaaatacttgaaaccccaaaaagcctctttgtttcttctgta-aagaaaaccaacaattgtgcataacaa
velvet worm         -gcaatatgcta-gacggaggggggataatttatgcaaatga------------agtaggcgctctgga-aaaatactttaaa-cctgaaaagcctctttgtgccttatgta-aagaaaaccaacaattgtgaataacaa
cactus worm         ----------tg--gctagagcccgataatgtatgcaaaata---------ttcgagacactctctaga-aaaatagtttaat-cgtgaaaagctcttttgtgcaaagcgtc-aaagcagccctcagtgttgaacaacaa
chiton              -------------------------ataatgtatgcgaatgtt--------ctcggcacacttatagca-ctgatacatggatccccaaaaggtctttttctccgttccata-tagaaagttaagaaacgtcgacaacaa
abalone             -----------------------aggcaatgtatgcaaatga-------------gtgtgcacccttagaacactgcccaaga-tttccacagattctttgt-atttctagc-aaagaaaccaacaatttgcagcgacaa
nautilus            --------------------------------atgcaaatga-----------gaaaaaacgcgcttcg-caactactctaaa-gctgaaaagattctttgtacggcctgca-aagaaaaccaacagtcgtgaataacaa
octopus             ------------------------------tcatgcaaatga-----------gaaaatgctggcttga-aaactacttttgg-cctgaaaagcttcattgtacacggtgta-aagaaaaccaacaatcgaaaataacaa
peanut worm         ------------------------------------------------------------------------------------------agaaagcctttgtgcctcccca-aagaaactcaataaatgcgaataacaa
shamisen shell      -----------------------------------------------------------------------------------------aaagaaccttttgtatgttagta-aggaacaccagcctcgttgaatgacaa
horseshoe worm      ----------------------------------------------------------------------------------------aaagaagtctttgtgtcctatgta-aagaaaaccaacaattgaaaataacaa
bootlace worm       --caacctgctc--tctctgcccagataatttatgtaaatga------------acggggcgcgctgga-aatatactttaaa-catgaaaagctgcattgtgttttgtgta-aagaaaaccaacaattgtgaataacaa

                        Sox_____
                    Hmx_______________
                                                         E-box_
                    ------
human GSX1          ag--aacaattaat-aagccgtcttttcaca----aaccagcagctggtttccccgtcaaggaaagttggaaaaaattcaggctgaatgcgcgcagaagctcttt------------
human GSX2          ag--cacaatt----aagctttctattcagg---agttctgcagctggtagcccat--tgggaaagtgggaggaaa-----------------------------------------
chimaera gsx1       ag--aacaattaat-aagccgtcttttcaca----aaccagcagctggtttgccgt--caggaaagttggaaaaaattcaggctgaatgcgcgcaaagctcttt-------------
chimaera gsx2       ag--aacaattaat-aagccgtcttttcacg----agctagcagctggtaacccgt--ggggaaagtaggggaaaatcagagatgaatgcgcgcaaaagtt----------------
lancelet            aaggggcgattgat-tacttggtgggtttcc------acggcagatggttaagg-----gggagaggtggcgggaggg---------------------------------------
acorn worm          ag--aacaattaat-aagttgttttttcacc----aaacggcagatggttcgaacg--agaagaagggagcgaaaagttgaaatgcctcgctggaaacaaaatgcgcgctttttaca
velvet worm         ag--aacaattaat-aagttgtttttctacc----aaacggcagatggttcgagcg--actgtaacatactcaaaagttgatatgcttttgcgaaactttt-tgcgctcctttttaca
cactus worm         ag--aacaattcac-aagttgtttttccacc----aaacgccagctggttaggccg--ctcgtaacatattcaaacgtcgagaggcgctgacgtaaccgcgatgc------------
chiton              ag--actaattaataaaattgtttttctacc----aaccggcagatggtttggacg--acggga-----------------------------------------------------
abalone             ag-aaacaattgcc-atgttgtt------------------------------------------------------------------------------------------------
nautilus            ag--aacaattaat-aagttgtttttctacc----aaacggcagatggctgg-----------------------------------------------------------------
octopus             ag--aacaattaat-aagttgtttttctacc----aaacggcagatggctgg-----------------------------------------------------------------
peanut worm         ag--acctgttaat-aagttggttttttgccttcaatacggcagatgttt-------------------------------------------------------------------
shamisen shell      ag--aacaattaat-aacctttttttccgca----aaacggcagatgttttctcg--------------------------------------------------------------
horseshoe worm      ag--aacaattaatcagaaagttctttcgtc----atacggcagatggt--------------------------------------------------------------------
bootlace worm       ag--aacaattaat-accttgcttttctggc----aaacggcagatgtccaacccg--atgggaacatatttaaaacttg-------------------------------------
```

**Figure 1:** *Gsx* enhancer

```
                                                                                                                                             CCAAT
human HMX3          ------------------------------------------------------------------------------------------ttcgcctgcctaatcccatttgccattgtacgcgcccaatcgccgt-ata
chimaera hmx1       ------------------------------------------------------------------------------------------ttttcttgagtaatcctatttgccattgtgcccgtccaatcgtgga-ata
chimaera hmx3       --------------------------------------------------------------------------------------ctctcccccttcagtaatcccatttgccattgtgcgtacccaatcgccgt-ata
lancelet            aaccggttgttgggatcgcccaataagaggggttttccaccactcgtccccattgttcaagatttgaga-ggtgtggtcacttgagcttttcgttcttcgtaatccaattacccattgtgtgtgtccaatcaccga-gtc
acorn worm          ---------------tcgcccaataagggtatgttttcaccactctgt-gcattgttatccatataagaaggcgtggttacagtggagtcgctttcactgtaatcctattgcgtcgtccccgtggccaatcacaga-gct
abalone             -------------------------------------------------cgatcgtcgatgattggagaggcgtggtgtactggaggattcctctttcattaatccgattagcttttgtgtttggccaatcagtgatttt
shamisen shell      --------------------------------------------------------------------------------------------------tctaatcctattatgtattgtcctcgaccaatcgc-ga-atg
stony coral         ---------------------------------------------------------------------------------------------tttctcatagttccattagccgtcattttagaccaatcacagc-agc
lace coral          --------------------------------------------------------------------------------tgattattcgagctttctcgtggttccattagtcgtcaccctggaccaatcacgga-cgc
starlet sea anemone -------------------ccaatcacagtggg-ttttatcagttt---tcccggttgtttgcaaagaagggggcggggaagtgcattttagctttctcacggtcccattagatgtattgctcgtccaatcagcgc-tct
pale anemone        ----------------------------------------------------------------------------------tgcattttctctttctcacggttccattagacacaaaggtcgaccaatcatagt-aca

                                                                EVX_______
                                   BARHL_____   NR4A______     POU6______
human HMX3          ctcccc---------gcatttaacttggatgacattttgatt--tcatcattagcatc---------------cggcgccggattgacgc---
chimaera hmx1       ctctttgcttgttaagcatttaacttgcatgacatttttatt--tcatcattatcatttttccccttct-----------------------
chimaera hmx3       ctttct---------gcattgaacttgtgtgacatttttatt--tcatcgttatcacc---------------cggcgccggattgacgc---
lancelet            gttttcagttgtcaagcatttaacttgtatgacacgtttatt--tagacgttattattgttatctctgctggacggcgccatttcgactcagc
acorn worm          tcattagcttgtcacccatttaacttgcacgccacttgtatt--tatatgttattagcgtg---ttagatgagcggcgccaatacgactc---
abalone             gcttgaagttgtcactcgtttaa----------------------------------------------------------------------
shamisen shell      cagtccatttgtcatccatttaacttctctgctactcttttctcttgtagttattatcgct---ccctgtgatcggcgccatcacggttaagc
stony coral         gcatttgcatgtcagacgtttaacaagtttgacatttttattcgtgatcattatcaccggc------ttaaattgtcgcctcgcatgc-----
lace coral          gcatttacatgtcagacgtttaacaagtttgacatttttattcttgatcattatcact-----------------------------------
starlet sea anemone ccgtttgcttgtcaaccgtttaacaagtttgacgtttttattcgcgatcattatcacaggc------ttaaaacgtcgccacttaagctca--
pale anemone        gttcttccttgtcaaacgcttaacaagtttgacatttttattcgcgatcattatcatcgac------ttaaaacgtcgccactt---------
```

**Figure 2:** *Hmx* enhancer

```
                                                                MEIS___    NFI______________                                      FOXO___
human MEIS1     -cagagcggatgatcattcattcaccacgttgacaacctcgcctgtgattgacagctggagtggcagaaagccatgagatttg-----gtagttgg---gtctgaggggcgc---tcttttttttccttt---tctttct
human MEIS2     tcatcatcaaccatcattcattcactaccttgacattccgggctttgattgacagctggagtggcaaaaagccatgaaacacg-----acagttcg---gttacatgtgggc---tgctgacgggccgct---cgtaacc
chimaera meis2  --acggcagaacatcattcattcaataccttgacaaccc-tgctctgattgacagctggagtggcaaaaagccatgagagacg-----acagtttt---gttacaagcgggc---tgttgacgggccgat---cgtaacc
lancelet        tcatgacagatcatcagtcattcagtactatgacaaccg-cgctttgattgacagcttgagtggcaaaaagccacgagacacg-----acagtttt---gttaca-ggcgct---tgttgacgggccgacattcatttcc
acorn worm      ---tgacatctcatcactcacgtccttccatgacaacct-cactttgattgacagcttgaatggcaaaaagccacgagacacg-----acagtttt---gttacagggctgt---tgttgatgggccgacataggtttcc
sea hare        ---tgacagctcatttattatccgggccctcgcggacggctcgtttgattgacagatttagtatca-gcatccacgagtcgcg-----acaggccctgttgttcgagtgtgctcgtgttgacgggc--------------
abalone         ---tgacagctcatttgttatcccgttctatagtaaccc-tggcttgattgacagatttagtggta--------------------------------------------------------------------------
oyster          ---tgacagctcgtacattatttcttgctagagtaacgt-ggttttgattgacatatt--gcgagaaatagccatgagttaag-----acaggtt---------------------------------------------
shamisen shell  --atgacagctcatttgttactcgtcactatggcaacgg--cggatgattgacagcctcgctgg-aagaggccacgagcatcg-----acagttta---gtt--------------------------------------
carnation coral ------------------------cctctatggaaactt-tagtttgattgacaggagatatggcaggcagccatgatacaaaccttgacaattc----gttacgact-----tgtgatgacgggggaatatgctatggg
stony coral     ---------------------------------------------tgattgacagacttgt-ggccaaaagccacgagacagg-cttgacaattc----gttactagt-----tgtgttgacggccgtacacatcttttc
lace coral      ---------------------------------------------tgattgacagacttgt-ggccaaaagccacgagacagg-cttgacaattc----gttactagt-----tgtgttgacggccgtacacatcttttc
pale anemone    ---------------------------------------------tgattgacagatcagtggggcaagagccacgagacaag-cttgacaatcc----gttacgggt-----tgtgttgacgggaatacacatctttct

human MEIS1     ttc------------------tttctttttttttttttaaactgattttt-------------------
human MEIS2     ttcagttcgggggcttgacaatttttttcttctttttcttctttttcttttcttt--------------
chimaera meis2  ttcagt-------------aatttttttttttttttttggcttgacatttttttt--------------
lancelet        ttctgt-------------tgcctgacattcttgtgtaagcttagaaagtccccggtacgttcaaactg
acorn worm      ttctgt-------------tagctgacacatttctgtaagcttagaaagtccccggcctgttcaaattg
sea hare        ---------------------------------------------------------------------
abalone         ---------------------------------------------------------------------
oyster          ---------------------------------------------------------------------
shamisen shell  ---------------------------------------------------------------------
carnation coral ttctcatt-----------tggttgacgggt---tgtaagcttagagagcc------------------
stony coral     cttacatt-----------gagttgacgggt---tgtaagtttagaaagctatccaactgt--------
lace coral      cttacata-----------cagttgacgggt---tgtaagtttagaaagctatccaactgt--------
pale anemone    tctgca--------------acttgacgtgt---tgtaagtttagaaag--------------------
```

**Figure 3:** *Meis* promoter

```
                                                    WRE__                       E2F_________               Smad_____
human MSX1       ----------------cccgtgcgccgggtgcagcctttgat-ccggccg-----------ccgctggcgccttaaaa--caacat-cctagtctgcctatt-----aggcgcgctgaggaatcgtgacacattgtttat
chimaera msx1    ----------------ctctccctgaagtcgcctcctttgat-cgcgccg-----------ctgctggcgcctaaaaa--caacataacttgtctgcctatt-----aaaggcactgactaatcgtgacagattgcttat
chimaera msx2    ----------------ccccgggggcgggggggggctttgat-caggccg-----------atcctggcgcctatgaa--caaccccacttgtctgcctctt-----aacttgtctgactaatggtgacagactgtttat
chimaera msx3    --------------tctccctgaggtagcgtctcggtttgat-cccaccg-----------aagctggcgccttccaa---aacataacttgtctgcctatt-----aaatgccttgacttatcttgaagaattgc----
lancelet         gggtaagtgttacatattcggagtgattgtggtccctttgat-cccgccg-----------ttcgtggcgcctagaaa--caacacaacttgtctgcgcatt-----gaaatatctgacaaatcaggtcagattgtttat
acorn worm       gggtaactgttccatatgctaagcaatgcatttctctttgat-cgggccgttaacgctctcttagtggcgcctgtacactcactaaaggatgtctgcttttttttaaaaaaaggatggcaagtcgtgatggattatttgt
sea urchin       --------------------------------------------------------------ttgtggcgccgactgc--cgtcctaagatgtctactcagt-----gaataaggtgacaaatggtgacggagtgtttgt
Satsuma tubeworm ----------------------------tggtctcctttgaa-gcgtccg-----------acgttggcgcctagcgc--gaacataacttgtctgtctatt-----cttcggcctgacaaatcgtgacagattgtttag
shamisen shell   ----------------ctatgcgcggggctgttttctttgatgtaccccg-----------accctggcgccttcttg--aaacttaacttgtctccctata-----gaaaagagtgacaagacttgactgcctgcttgc

                     MSX_____
human MSX1       ccccgcaattagcgatg------------------------------------------------------------------------
chimaera msx1    agtcgcaattagcaatgcagacgctc-ggcgg---------------------------------------------------------
chimaera msx2    ccgcgcggttacccagcaggtgccgc-gggg----------------------------------------------------------
chimaera msx3    -----------------------------------------------------------------------------------------
lancelet         cctcacaattagtactggctaggtca-caaggtaattgatcactc--------------------------------------------
acorn worm       cgagccaattagctctggcttggtcc-gagggtaattgatcacct----aaaggggtgtgtactcattgcaaataactcctttatcttc
sea urchin       acttgcaattagcctggctatgcctcctgaggtaattgatcgcccgaatagaaagagccgttcccactgaaaaaaccctctccctcctc
Satsuma tubeworm acgcacaattagcgacgccctgacgc-gg------------------------------------------------------------
shamisen shell   tcaagcaattag-----------------------------------------------------------------------------
```

**Figure 4:** *Msx* enhancer

```
                                                                                              E-box_ FOX_______                         TEAD____
human SIX1     gtgcgcacttgctccagactttagggtattaactgatttttagagccgggtttaagcctccgttc---cggtcctcacacagctgccatgtcaata--agga-ctgtttcaagca---gaggcattccttctgggagggc
chimaera six1  gtgcgcgttcgtttggtgatttacggtacaatctgactccgatag--agggttaaggatcagttc---cggtccccaaacagctgccttgtcaata--aggatctgccacaagtg---gagacatccctgctgtcaggac
lancelet       -------------------taccttatacttactaatactgagtc--cgatccacgtattggttc---cggtatcgaaacagctgttttgtcaata--agaa-ctgtcaaacgaa---aaagcattccaacgatgccggt
velvet worm    ----------------------------------------------------------------------------------------------------------------ggg---gaagcattccgctaataccagt
shamisen shell -------------------taccttacactgatcaatttagagcaaaacgttagagcattggtccaagtagcactggaacagctgttttgtcaatattagcg-cagtctggggagcataaagcataccactaataccggc

                           MEIS___
human SIX1     --------------------------------------
chimaera six1  --------------------------------------
lancelet       gatgtactctacctgtcaaacattatctctaataacat
velvet worm    gatgtact-tacctgtcaaatattatctttacaaacat
shamisen shell -----------cctgtcaaatattatct----------
```

**Figure 5:** *Six1* enhancer

```
                                     PKNOX__________
                                  GFI1________
                         Six________          RFX____________                                                               CCAAT
human SIX2        --------------cccgccgaggctcgggttaccagtgactgacagcgtctccatggcgaataatttgactccgactattgtctggcgctggcaggcccc---gggtcagataaccggaccaatcagggcgcgggccgc
chimaera six2     -------------------------tcgggttactctgattgacagcgtctccatagcaaataatttgact-taactattgtcttatcctcaccatccgt----aggtcagatctccgaccaatcgtatgctacacaat
lancelet          atactcttttgacacctaagtgcatctcatgttacattgattgacattgtcgccatagtgaataatttagt--aaaccattggaataatatcgatatatcc--ggcgcgcacactcctgaccaatcacggcaaagcagct
acorn worm        acgttctgttgacacctcatcggcgatcaagtttcactgattgacgtggttgccatggcaaataattcgac--aaactattgtatgaatatctatatatccgtgg---cgagcaagcggaccaatcagggcagagtatct
velvet worm       ----------gacacctaaagcagggtagggtttcattgattgacggcgttgtcatgacaactaattttgt--caagccttctatgaatattgatatattggtggggtcggaagaaccaaccaatcagaagcgtgaataa
abalone           -------gtccacgcctcttgatgggtcgggtttca-tgattgacattgttgctatggcgatggattttca--caggcattgtgtgaataacgatatatccatggggtcgcgcgccagagccaatcagcggccagcagag
owl limpet        ------------cgcctcctgccctatcaggtttcagtgattgacgtcgttgctatagtaatagattttca--caagcattgtgtgaataacgatatatccacaggg--------------------------------
oyster            ------------------------------------ttgattgacgtcgttgctgtggttactaatttctc--acggcactgagagaataagcgatatcttctggggtcgccagaaacagccaatcatctgaaagtagtg
peanut worm       ------------cacctctcagcgtatcaggttacagtgattgacgtggttgccatggtgactaatttctt--gcagcattgtatgaatatcgatatagccggcgggtcgctgcgcacggccaatcacggcccggatgcc
sandworm          ------------------------tgtcgggtttcagtgattgacggcgtttccatggagaataatttggt--agagtattgcatgaatatcgatatgtccacggggtcgaggccatgggccaatcagcgtgcagccggc
Satsuma tubeworm  -------------------------atcgggtttccgcgatagacggtgttgccatggcaagtaatttgtc---gcgcgttgcatgaatatcgatatgtc---------------------------------------
shamisen shell    ------tatcaacaccccttcccgtatcaggtttcactgattgacgtggttgccatggtgtaccaactcac--taagtattgtatgaatatcgatattagcaaggggtcgggaagctgaaccaatcagcttctagcttac

human SIX2        cgcgcctcatgcccgcttag-aataatattattaaga---------------------------------------------------------------------
chimaera six2     gactattcgtgccaacttgg-aataatattattggaaaaaaaagaaacccccca----------------------------------------------------
lancelet          ggctattaatgtcgccctaa-aataataatgtcaaaaagtag--caaccccacacgttggcctgtttgtagtgtcgtttttttccctgctacacacacacacgcacg
acorn worm        ggctattaatgtcgcgttaa-aataataatgtcagaacgcgg--caaccccacacgttgggttatttgcactgttgcttctca-------------------------
velvet worm       aactattactgtcgctttaa-aataataatgtcaaagcatcc--cagccccacacgttggcttgttttcaagcttatctttcaccccgaagacgaatattcaagcagg
abalone           ggcca-------------------------------------------------------------------------------------------------------
owl limpet        ------------------------------------------------------------------------------------------------------------
oyster            aaccattactgcacctcta-----------------------------------------------------------------------------------------
peanut worm       gcctattactatcgccttga-aataataacgtcaagatgttg--gtctctcacatgttaactctgttt----------------------------------------
sandworm          ctcctgtaacgttgctttgaaaataataaggtagaggcagg----agcccc---------------------------------------------------------
Satsuma tubeworm  ------------------------------------------------------------------------------------------------------------
shamisen shell    ttccactaatgtcgccttaa-aataataacgccaagatgtg----agccccacacgttgggttgttttcac-------------------------------------
```

**Figure 6:** *Six2* promoter

```
                                                                  Nrf1___________          Yy1______                HOX_______
human ZFHX3    tattgatccagaaggtggctaattagccctt--cccgtcgtgcctgggtaatg-aagcacatgcgcactgaattaatcacagccattttttt------------gcaggaaactaattagcagaataaagatccaaagact
human ZFHX4    ---------------------------------------------------aaagacatgcgcactcaacttaat-cagccatttttttaaacagggctaaaacgataataattagcagaataaagacatatcggat
chimaera zfhx3 tattgatccgagggttcgctaattagccctttacccgtcgtgccttcgtaatg-aagcacatgcgcactgaattaatcacagccattttttg------------cacggaaactaattagcagaataaagatccatcgact
chimaera zfhx4 --------------tcactaattacaccttt--cccgtcgtgcatcggtaatgaagacacatgcgcactcaacttaat-cagccatttttttttacaggggc-------aaaataattagcagaataaagatatatcggat
lancelet       ----------------------tagcctat--cccgtcatgccc--atcgcg-acagacatgcgcgctgcgctaat--cagccattttac-------------acggaaaataattagcagaataaa---atatcgaat
acorn worm     ------------agatgtaaattgtctttc-gctta-----------------ttacgcatgcccattga-tgcctcccagccatttt-------------------ggaataattagcaaaataata--atatcgaat
sea urchin     ------------agattgggattaccttgc-accta-----------------ttacgcatgctcattgattgcgctcctgccattttac--------------acaaaaataattagcaaaacaat------------
abalone        ---------------------------------------------------------------------------------------------------------taattagcgcaataaa---atatccact

human ZFHX3    ttttt------------------------------------------
human ZFHX4    tttcatttc--ctttcctccttttccc--------------------
chimaera zfhx3 tttttcatt--gcgagtttcgccctcttttccgcc-----gcgatcgc
chimaera zfhx4 tttcatttc--cagtttccccttttccgttt---------gcggtcgc
lancelet       tttcatttg--gttatttctagtttctttctcggcgccgagcgcttct
acorn worm     tttcatttg--gttatttttagctctttatttttcacctagcatttgt
sea urchin     ------------------------------------------------
abalone        tttcatttactgttatttttagctgccgttttg---------------
```

**Figure 7:** *Zfhx3/4* promoter

```
                                                       NRRE__       NRRE__                      POU4__________
human EBF1     ---------------------ttttacttcctgtttcaa-----aggtcaaggcaggctgacctcaacagcagtgtggaggctgctgaattattcatatcattgactttgtgtca-----gcttccacatttgtggaga
human EBF2     ttctgtggtcactattattttcttggcctacccatcacaaaacagaggtcacagcaccatgaccttttcctggtggaatggactgcttaagaagtcacatggccaagtttctatcg----ggggtccacatttgtttcct
human EBF3     --------------tgtttcctttcacttcctgtcacaa-----aggtcaaagagagccgacctttgc-tggcagtcctgctcactgaattattcatgggattgactttttgtca-----gcacacacagttgtgctct
chimaera ebf1  ----------------ttcccctccacttcctgtcataa-----aggtcgaggcagggtgacccctgcatcagcagcaaggttgctgaattattcatgcaattgactttgtgtca-----gcgtacacatttgtgcaaa
chimaera ebf2  ------------------------------------------------------------------------------------------------------------------------------cacatttgttcttg
chimaera ebf3  tttcttagtcaacattgcttcctttcacttcctgtcacaa-----aggtcaaagaaagctgacctttgc-tggcagcttcagttgttgaataattcatgtgattgactttttgtca-----gtgcacacagttgtgctct
lancelet       tg-------------tctcctaactcatttcttacttgcg-----gggtcaggctctactgaccttt---agcggcaccagtaccatgaattatttataaggcaaaattattagaa-----atgtacacagttgtgtttg
velvet worm    -------------------------------------------------------------------------------cattaattattaatgaagaagcgttatgattagtgtggtacacacaactgcggatt
chiton         -------------------------------------------------------------------------------------------------------------------------gtacacaggtgtcctct

                   ZNF24________                    E-box_
human EBF1     gggatgttttattcatttatctcatttagggttcttctctgcacctgtttctaccattaagtaccctgtaatttgcatttgatgctaatttagtt-tcatatcagcctggtcccccttgttaa-tttaatttt------
human EBF2     ccactcttttaatcagaatgctcatttaattct-attccagcacctgttc--agcaattaacaccctacgatttgcatcttaccctga-----------catttgctatctttccaa---ttagtttttatatcagtata
human EBF3     gggacattttattcatttacctcatttaaagcgcctttctgcacctgttc--aagtattaatatcgtgtaatttgggcctaatgccgattttgct-gaacactcattatatttttta---ttag-ctatatttccaaacg
chimaera ebf1  gggatgttttattcatttatctcatttaagggcctttcctgcacctgttcttaccactaagtacccagtgatttgcatttgatgctaatttagttatcgtatcggactggttttcctc--ttaa-tttcatttt------
chimaera ebf2  agaccgctctaatcacactcctcatttagacct-acacccgcacctgttc--agcaattaacatcacagattttgcatttaacat-----------------tttgctgtgtttccaa---ttag-ttttatcgctcagta
chimaera ebf3  gggacattttattcatttacctcatttaaagcgcctttctgcacctgttc--aactattaatatcatgtaatttgtgcctaatgccgattttgct-taacattcattatatttctta---ttag-ctatatttccaaatg
lancelet       gggcacattggctagcccgttccatttgggtg--ttctctgcacctgtca--tgcaattagtgcgttataatgttcctgttgtgcccaatttggt--ggtgctcattatcttctgtaat-ttgg-attcattatgaaatg
velvet worm    gggagacttcattcatttaactcatttaaaggg-ttttccgcacctgtca--gatcataaataagccctaattttcacttaatgccaatttagct-taacagtcattattttctata---tcag-gt------------
chiton         gtgagatttgattcattttctccattttgcgcgcctttcaacacctgttc--aggggtaaata--------------------------------------------------------------------------

human EBF1     -------------------------------------------------------------------------------------------------------------------
human EBF2     attaggtagggttattacagtgcatcaagcagggtcat-ggttattcccaagcctttaagcgctttccca-gatttgtttagggaagccactaagggcaaatggttga
human EBF3     attatgtatgattattac-------------------------------------------------------------------------------------------------
chimaera ebf1  -------------------------------------------------------------------------------------------------------------------
chimaera ebf2  attagacgaggttattaccacatattaaactcggcaacaaagaattcccacggctttaatctttctctc-----------------------------------------------
chimaera ebf3  attatgtatgattattaccagaccctaggaacagtgatgggttattccctagacctttacatcttcttgtcaggctgttttcggaag-cagtgagaataaacatttga
lancelet       atcatacaggattgtgac--------------------------------------------------------------------------------------------------
velvet worm    -------------------------------------------------------------------------------------------------------------------
chiton         -------------------------------------------------------------------------------------------------------------------
```

**Figure 8:** *Coe* intron element

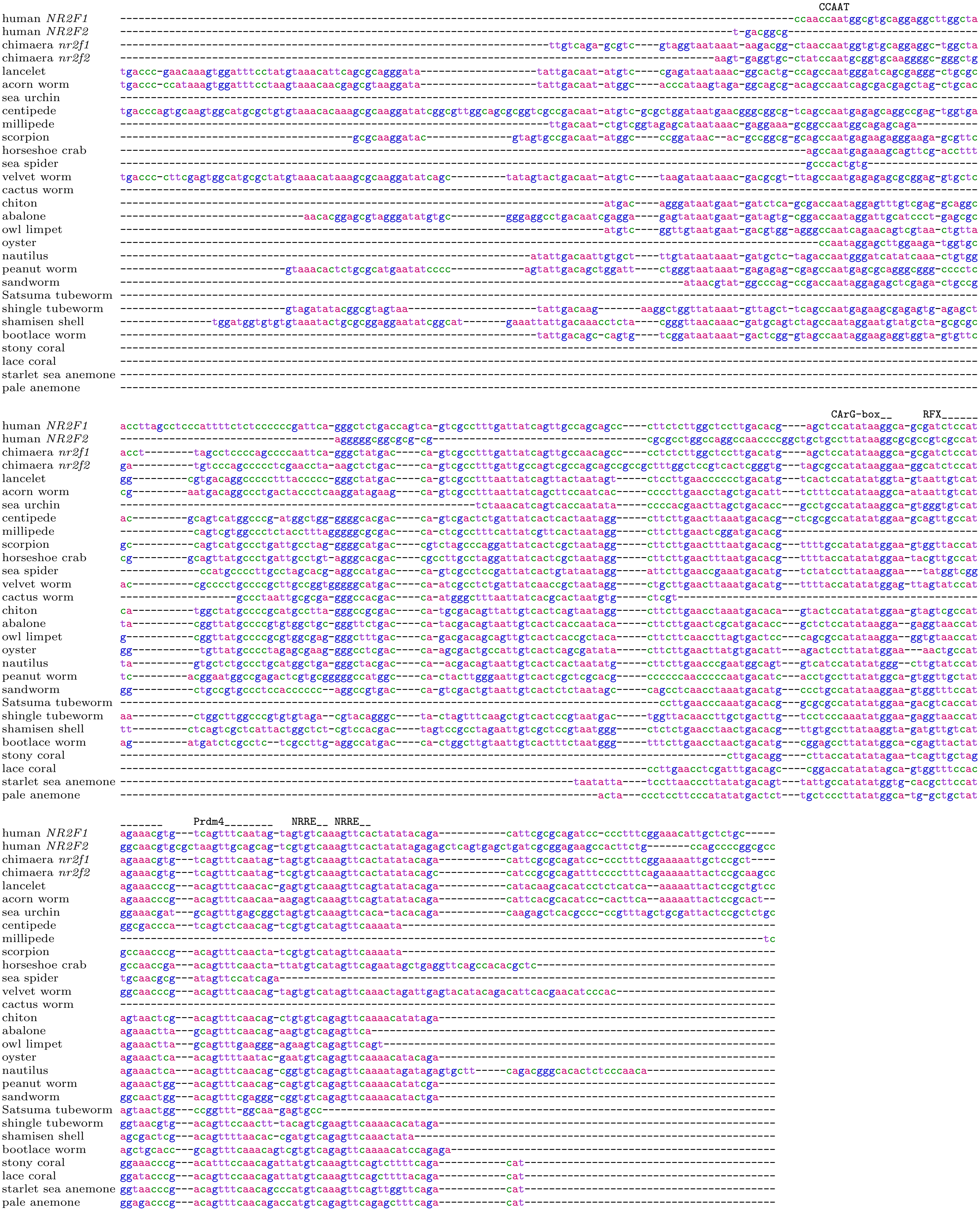


**Figure 9:** *Nr2f* promoter

```
                                                                                                                                   RFX______________________
                                                                          RFX_____________________              RFX__________________  E2F_________________
human RFX1         ------------------cgcccccgcgcgtgacgtaactagggtgggtagcaacagttgccccggtgagggaaacggaggcgccatagccacggtagtcg-t----ggcgaccaagca-accc-----ggcaacgcg--
human RFX3         ----------------------------------------------tatagcaaccgttgccttgtgaat-------caagcgccatagtcaccgtagtcc-t----ggcgact--gtt-accc-----atcaacaacaa
chimaera rfx1      ------------------cgccgcgttgcctgggttgattccggaggctagcaactgttgccttgaggac--------gagcgccatagccaccgtagtcc-t----ggcgact--gtt-accccgtcaaacaacaactt
chimaera rfx3      -acgg-----------------------------ttgctgggctcttatagcaactgttgccttgtgact--------cagcgccatagtcaccgtagtcc-t----ggcgact--gtt-accc-----accaacaacag
lancelet           ggcgggtaggcggagt-gatccgtgttgtgtgtgcgtgtgtcgccgtgtggcaacagttgccttggttgt--------gtgcgccatagccaccgtagtca-c----ggcgact--gtt-tccc-----gccaacaacac
acorn worm         ---------------------------gtgtactgtggttaggttatttggcaactgttgccttgagaat--------aaacgccatagccaccgtagtcc-t----ggcgact--gtt-accc-----accaacacggc
sea urchin         ---------------------------------tacagtggtgaggtttagcaacagttgccttgagaat--------aaacgccatagtcaccgtagtcc-t----ggcgact--gtt-accc-----gccaacga---
centipede          ------------------------------------------------tggcaactgttgccttaggaat-------ccgacgccatagacacggtagctg-t----gacgactgcgta-acccc----atcaacaggac
millipede          ------------------------------------------------tggcaactgttgccctaggaat--------aagcgccatagacacggtaactg-t----gacgact--ggt-agcc-----atcaacagtaa
tick               -------------------------------------gttgcggcagttggcaactgttgcccttggagt--------caacgccataactacggaagctg-c----aacggcttggta-acct-----gtcaaca----
horseshoe crab     -----------------------------------------------ttggcaactgttgcccttggaat--------caacgccataccgacggtaaccc-g----gacaacc--aaa-atct-----gtcaaca----
velvet worm        ------------------------------------tgtacagaggtttggcaacagttgccttaggaat--------aaacgccatagccaccgtagctg-t----gacgact--gtt-accc-----gccaacga---
cactus worm        ----ggtcggcgcaataatctattgccctgcatcgatgtgatccggtttggcaactgttgcccaagg-----------aaacgccatagacacggtagctg-t----ggcgact--gtt-accccg---aacaacaaagc
chiton             ggcggttatgcggaataat----cgttgatttgctgacgcgctgagtctggcaacagttgccgtaggaat--------aaacgccatagccacggtagctg-t----gacgact--gtt-accc-----gccaacaagac
sea hare           -------------------------------------------gagtctggcaactgttgccgtaggaat--------aaacgccatagacacggtagctg-t----gacgact--gtt-tccc-----atcaacaacta
abalone            -------------------------------------------gaggctggcaacagttgccgtaggaat--------aaacgccatagccaccgtagctg-t----gacgact--gtt-accc-----accaacacgac
owl limpet         -------------------------------------------gagtctggcaacagttgccgtaggaat--------aaacgccatagccaccgtagctg-t----gacgact--gtt-accc-----accaacagaac
oyster             ------------------------------------------ggagactggcaactgttgccgtaggaat--------aaacgccatagccaccgtagctgtt----gacgact--gtt-accc-----accaacacgta
nautilus           ----------------------------------------------tctggcaactgttgccgtaggaat--------aaacgccatagccacggtagttg-ttgacgacgagt--gtt-accc-----gccaacaaga-
octopus            -------------------------------------------gagtgtggcaactgttgccctaggaat--------aaacgccatagccacggtatctact----gacgact--gca-gccc-----gccaacaacac
solenogaster       --------------------------------------------agtctggcaactgttgccctaggaat--------aaacgccatagacacggtagctg-t----gacgact--gtt-accc-----accaacaactc
peanut worm        -------------------------------------------gagtctggcaacagttgccgtaggaat--------aaacgccatagccaccgtagctg-t----gacgact--gtt-accc-----gccaacac---
sandworm           --------------------------------------------cgtctggcaacagttgccgtaggaat--------aaacgccatagccacggtagctg-t----gacaact--gta-tccc-----gccaacaagac
Satsuma tubeworm   --------------------------------------------cttctggcaacagttgccgtaggaat--------aaacgccatagccacggtagctg-t----gacgact--gtt-accc-----gccaacaagac
shingle tubeworm   --------------------------------------------cgtctggcaacagttgccgtaggaat--------aaacgccatagccaccgtagctg-t----gacgact--gtt-accc-----gccaacacgac
shamisen shell     --------------------------------------------cgtctggcaacagttgccgtaggaat--------aaacgccatagccaccgtagctg-t----gacgact--gtt-accc-----gccaacaagtc
horseshoe worm     --------------------------------------------cgtctggcaacagttgccgtaggaat--------aaacgccatagccaccgtagctg-t----gacgact--gtt-accc-----gccaacaagaa
bootlace worm      -----------------------------------------cggagtctggcaacagttgccgtaggaat--------aaacgccatagacaccgtagctg-t----gacgact--gttgaccc-----gccaacaagac

human RFX1         -----------------------------------------------------------
human RFX3         ctac---tcctctcctcctcctcctcctcttcctcctcctcct------ccccaccacc
chimaera rfx1      acgccacaaaggtgattctttttctctctctctctcttctctc----------------
chimaera rfx3      gagc---gcaggtgatctcctcctcactct-----------------------------
lancelet           caacc-acaaggtgattttcctccatcctctactcccactaccgcaccgcgccactacc
acorn worm         a------taaggtgatttttttcctttcagttcctagtcttcc----------------
sea urchin         -------caaggtaattttttcctttttta------------------------------
centipede          ------agaaggtga--------------------------------------------
millipede          -------cgaggtgattctctc-------------------------------------
tick               -----------------------------------------------------------
horseshoe crab     -----------------------------------------------------------
velvet worm        -----------------------------------------------------------
cactus worm        cagc---tgaggtaatttttccttc----------------------------------
chiton             -------taaggtgatttttcctt-----------------------------------
sea hare           -------taaggtgaattt----------------------------------------
abalone            caac---taaggtgatttttcttttccttt-----------------------------
owl limpet         -------taaggtgatttttcttta----------------------------------
oyster             -------aaaggtgatttttc--------------------------------------
nautilus           -------gaaggtgatttttcttttccatttcggttcgtag------------------
octopus            acacc-aaaaggtgattttccttttccc-------------------------------
solenogaster       -------cgaggtgatatacttattattc------------------------------
peanut worm        --tc---taaggtgatttttc--------------------------------------
sandworm           -------taaggtgattttt---------------------------------------
Satsuma tubeworm   t--------aggtgatttttcctttac--------------------------------
shingle tubeworm   -----------------------------------------------------------
shamisen shell     -------taaggtgattttcct-------------------------------------
horseshoe worm     c------taaggtgattttt---------------------------------------
bootlace worm      -------taaggtgatttttacttcctattttgctctaactta----cacgcc------
```

**Figure 10:** *Rfx1/3* promoter

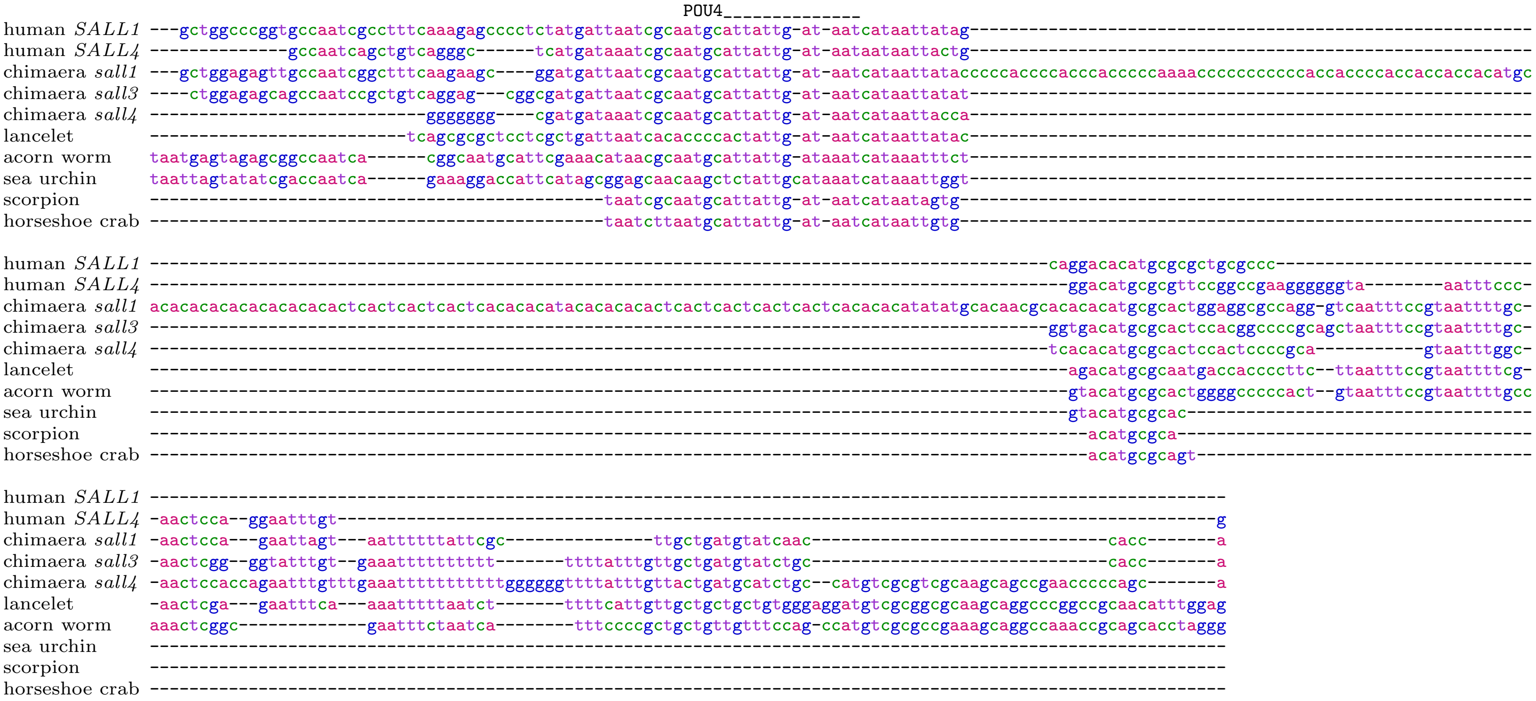


**Figure 11:** *Sall* promoter

```
                                                                                     CEBP__________                       E2F____________ Smad____    E-box_
human             ----------------------------aagagtt-----------aagaaacttgaa-accttgtttgcgcaacaatcag-------c----gcc---gcgg-agccgccaaagtgtctagac-tggcatatgatggga
chimaera          ----------------------------aagggtt-----------aagaaacttgag-accttgtttgcgcaaccatcgc-------tgcagatg---agag-ccgcgccaaaatgtctagaccggccacatgattgga
lancelet          ----------------------------------------------cggtaatgtgat-ctgttatttgcgcacccatcgg------------------cgga-gcgcgccaaaacgtctagac-gcgctgccgccgcac
acorn worm        actgtttttcggtgagatattc---cgggggagaagtaacagccaagtgtaatttgag-ctgttatttgcgcaacaatcat-------tttagact---cggg-gcgcgccaaaatgtctagac-gaccacgtgataatt
sea urchin        -------------------------------------------------------------gttatttgctcaacaatcat-------ttcagact---cgag-ccgcgccaaaatgtctagac-atccacatgacccaa
sea urchin        -------------------------------------------------------------gttatttgctcaacaatcat-------ttcagact---cgag-ccgcgccaaaatgtctagac-atccacatgacccaa
centipede         ----------------------------------------------cggtaattcgat-ttgttatttgcgcaacaatcac-------tgaagaca---tggg-gcgcgccaaaaagtctagacgaatcacgtgtccgct
millipede         attctttttcgcgaaaatggcgagaggagaagcatataatcagtagccgtaatttgat-attttatttgcgctacaatcgt-------ccgagacag-tgtgc-gcgcgccattgagtctagac-gatcacgtgat----
scorpion          -----------------------------------------------agtaatttgat-ctcttatttgcgcaacaatcgt-------ttcgg-----------gcgcgccgaaaagtctagac-gtccacgtgatcggt
tick              -------------------------------------------cgaaggtaatttgat-cggttatttgcgcaactatctcgcgcagacagaagga---ggag-gcgcgccgtcctgtctagac-gggcacgtg------
horseshoe crab    --------------------------------------------------------------ttatttgcgcaatgatcac-------cttaaaca---ccga-acgcgccaaaatgtctagac-actcgtgtgagcata
horseshoe crab    --------------------------gggaggaacgtagtctcagaaagtaacttgat-cacttatttgcgcaacaatcgc-------atcagaca---cta--ccgcgtcaaaatgtctagac-acgcacgtgaacata
horseshoe crab    -------------------------------------------------taatttgat-ctattatttgcgtaacaatcgt-------atctgtgc---ttag-ccgcgccaaaatgtctagac----------------
horseshoe crab    -----------------------------------------------agtaatttgag-ctcttatttgcgcaacaatcgc-------ttcagaca---cca--gcgcgccaaaatgtctagac-gtgtacgtaata---
velvet worm       --------------------------gggaggaatgtaatcattggaagtaatttgat-atgttatttgcgcaacaatcac-------tttagaca---tggg-gcgcgccaaaaagtctagac-aagcacgtgactaac
cactus worm       --------------------------------------------------------------------------cagtcac-------tatagatg---tgtg-gcgcgccagcgagtctagac-ggccacgtga-----
abalone           ----------------------------------------------cagtaatttgat-atgttatttgcgcaacaatcat-------agtcgaca---tcag-ccgcgccaaaaagtctagac-acccacgtgctacat
nautilus          ------------------------------------------------gtaatttgat-atgttatttgcgcaacaatcac-------tttaggca---tcagtccgcgccaaaaagtctagac-tgccacgtgaccgtc
octopus           -----------------------------------------------tgaaatttgatcccgatatttgtgcaacaatcact------gaaagacagagtgag-ccgcgccaaagtgtctagac-tagcacgtgacccgg
peanut worm       -----------------------------------------------agaaacttgat-atgttatttgtgcaacaatcac-------tgtagaca---gtgg-ccgcgccatggagtctagacgagccatgtgaccatt
sandworm          ------------------------------------------------gaaacttgat-acgatatttgcgcaacaatcac-------tttagaca---gcga-ccgcgccaaaaagtctagacaaggcatgtgactgaa
Satsuma tubeworm  --------------------------------------------------aactcgat-gccttattggcgcaacaataac-------tttagaca---gcta-ccgcgccaaaatgtctagacctcgcacgtgacccgc
shamisen shell    -----------------------------------------------agtaattcgat-aggttatttgcgcaacaatcac-------tttagaca---tcag-ccgcgccaaaaagtctagacaaaccatgtgatcaag
stony coral       ---------------------------------------------------attgaaa-ttggtatttacgtaacaccctc-------atcagacaactatag-acgcgcctggctgtctagac-aagcatgtgacagga
lace coral        ---------------------------------------------------attgaaa-ttgttatttgcgtaaca-ccac-------atcagacaactatag-acgcgccaatgcgtctagac-atgcacgtgacaata
```

```
                        CCAAT
human             ggcag-ccaatgactccgcggcgctcc---------------------------
chimaera          gccaa-ccaatgactcaaaagtgtccc---------------------------
lancelet          gga---ccaatcagcgcgcagct-------------------------------
acorn worm        ttgaa-ccaatcaagtgttagcaatgg-tgtgg-cagtaagtgttgtaaatttc
sea urchin        gtcaa-ccaatcagcgtagaagtaatcgtgtggtcagagagagtagtaaatttc
sea urchin        gtcaa-ccaatcagcgtagaagtaatcgtgtggtcagagagagtagtaaatttc
centipede         cccgaaccaatggcggcgc-----------------------------------
millipede         ------------------------------------------------------
scorpion          tatag-ccaacc-------------------------------------------
tick              ------------------------------------------------------
horseshoe crab    ctcag-ccaatggtt----------------------------------------
horseshoe crab    ttcaa-ccaatagc-----------------------------------------
horseshoe crab    ------------------------------------------------------
horseshoe crab    ------------------------------------------------------
velvet worm       ataaa-ccaatcagatct-------------------------------------
cactus worm       ------------------------------------------------------
abalone           cttgg-ccaatcagcgtgcagct--------------------------------
nautilus          gtcgg-ccaatca------------------------------------------
octopus           ccgcc-tcaaccaatcaga------------------------------------
peanut worm       tccga-ccaatca------------------------------------------
sandworm          attgg-ccaatggc-----------------------------------------
Satsuma tubeworm  cccag-ccaat--------------------------------------------
shamisen shell    ataag-ccaatca------------------------------------------
stony coral       atcaa-ccaatcaa-----------------------------------------
lace coral        tttca-gccaat-------------------------------------------
```

**Figure 12:** *Smad6* promoter. This matches four places in the horseshoe crab genome: three upstream of genes annotated as “smad6-like”, and one upstream of the edge of a genome fragment.

```
                                                                                                                          HOX___________
human             -------------------------------------------cttcaagt---agcccccattgaatagac--------taagttgacactgcgtgacagtgaaacaacataataaaaaatacatgagc--------cc
chimaera          -------------------------------------------cttcaagt---agcccttattgaatagac--------taagttgacactgcatgacagtgaaacaagataat-aaaaatacatgagc--------ca
lancelet          tgaatgggttcgccgctgtaggcctccattgttcccgcttgttctgcgtat---taattatacaaagggggc--------tgcagtgacacaagaggcaaaaataacaacataataaaaaatacatgagc--------cg
acorn worm        tgaatggaccggttactctatgcggccattgttccgccttgttc-gggcat---taatgttataaaagagtt--------tgctgtggcacaagaggcaaaaataacaacataataaaaaatacatgggc--------ct
sea urchin        -------------------------ctattgtccaagcttgttcagcgtat---taattatataaaggacccgtggtttggggaggtttgcagaacggaaaaataacaacacaatcaaaaatacatgaacgccttcaata
stony coral       ------------------------------------------------------taattatagaaaagaa-----------acagaaagacaaggaaaacgaataacaacacaataaaaaatacatgagg--------ag
lace coral        -------------cgctgtgcgtcgttattgtaaatcttactttcgcatattgataattatataaaagaa-----------actgagagacaagggccgagaataacaacataataaaaaatacatgagg--------tg
pale anemone      ------------------------------------------------------taattatataagagaa----------tataggtgaacaagaggcgagaataacaacataatcaaaaatacatgagg--------tg
```

```
                                MEIS___________
                                   FOX___________                           SOX_______                                        SOX_______
human             ctgaataggagcaggcgcataaataaataaaatgggtgaccaaaactggataaactgaatgacaaaacggtgaaaggggaacaaaaagatatttaacacgctagattagcattagaatgcgatctacaaggcaga--aca
chimaera          ctgaatagaagcaggcgcataaataaat-aaatgggtgaccaaaactggataaactgaataacaaaacggtgaaaggagaacaaaaagatatttaacacgccggattagcattagaatgcaatctacaaggcaga--aca
lancelet          ctgaatgcccgaactgacataaataaat-aaatgagcgatcaactgtagataaactgaataacaaagat-tgaatggcgaacaatgag---tataatgcgactgatctacatgacaaaagaatatacatactgga--aca
acorn worm        gtgaatagctgatatgacataaataaat-aaatgagcaattcacacaaagggtgttgaatagactagtt-tgaatgacatacaatagg---tgtaacgctgctaatttgcatgacaatagggcatacaacttagt--aca
sea urchin        gcgaagggtgcgcatgacataaataaat-aaatgag--------------------------------------------------------------------------------------------------------
stony coral       ttgaatgaaatagatgacataaataaat-aaatcgacgatcaagaagagataaaatgagtaacaaaagt--gagtgaattacaattgc---tctaa-actgac-atctccataacaatggattaca--------------
lace coral        ttgaatgaaagagatgacataaataaat-aaatggacgatcaagaagagataaaatgaataacaaaggt--gaatgaattacaattgc---tctaatactaat-atctccataacaaaagattacatccacaaaaataca
pale anemone      atggatgaaagcgatgacataaataaat-aaatagacgatcaagaggagataaagtgaataacaatagt-ggaatgaattacaattgt---cttaatactcct-atcaccataacaatgacatacatcaacaacgc-aca
```

```
human             attga-tgaatag-gtttaccggc--caagaaagaaatggactaaat-gccctttgaatagatatgcttt----------
chimaera          attga-tgaatag-gtttaccggc--cgagaaagaaatggactaaat-gccttctgaatggatgtgcttt----------
lancelet          aaggc-tgaatag-aacgaccggg--gtataaataaatagacccaaa-caggcttgagaagaatggcgtgaatagaaagg
acorn worm        aaggcgtgaaggg-atcgaggggga-gagtaaataaatagaggcaaa-cagcattgaattggtggctttgaatagagagg
sea urchin        --------------------------------------------------------------------------------
stony coral       --------------------------------------------------------------------------------
lace coral        ataac-tgaatag-------------------------------------------------------------------
pale anemone      ataag-tgaatagcggcgacagaaagatctctgtaaataaaccaaaaccaactacgataggaatgg--------------
```

**Figure 13:** *Sox21* enhancer

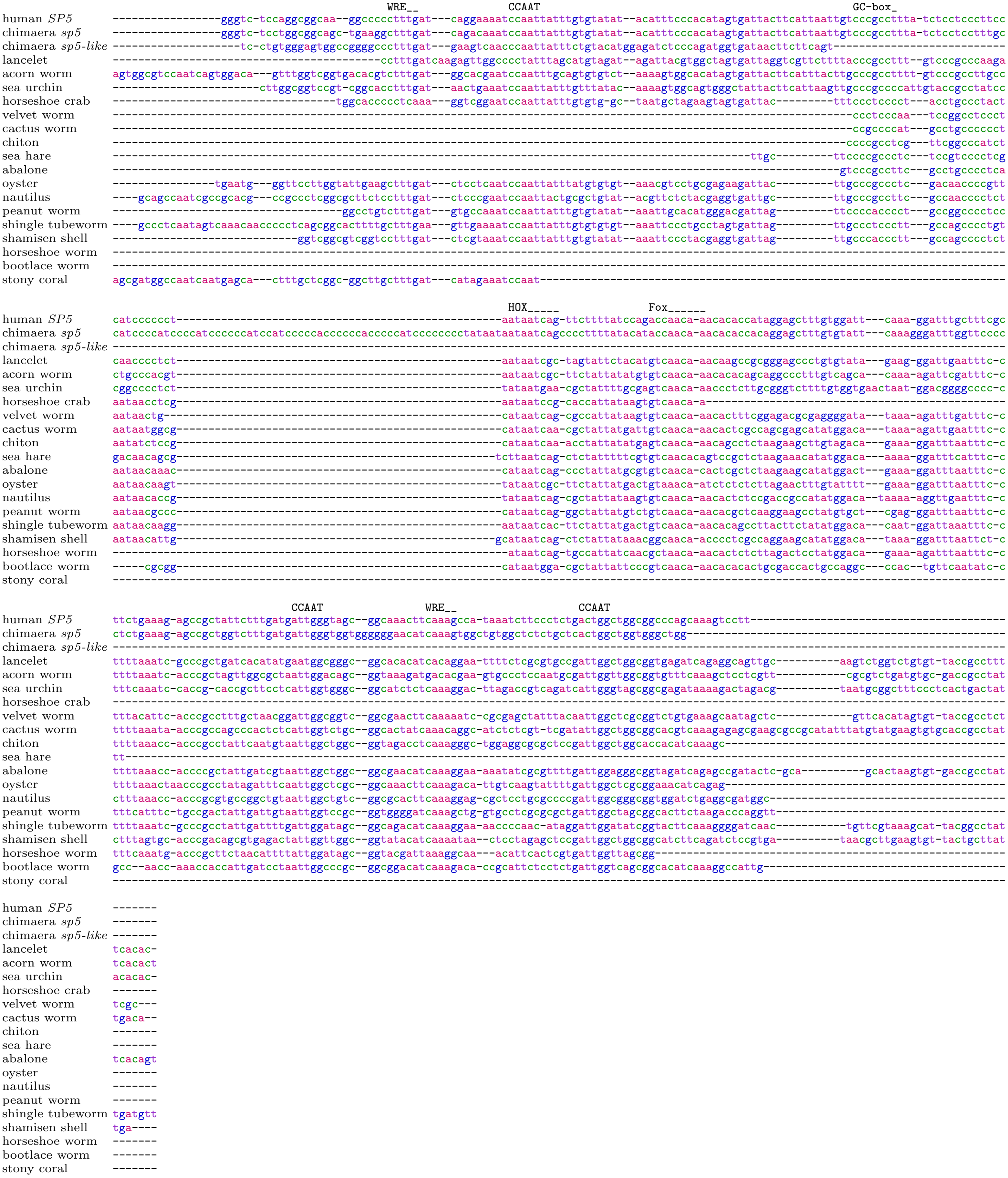


**Figure 14:** *Sp5* promoter

```
                                                                              FOX________
human TBX2     gggggccggggcaggaggaggggtctcggggcccggcggcccgtcattggttaatattttattctgttgacatgttttcttact------------------------------gctgaggcttccgacaccttct----
human TBX3     ----------------------------------------------------------tgatcatgttgacatattcacaggacaggcagtagtaccgatgcggc-gctgcgacgttacagtttccgacaccttctttt-
chimaera tbx2  gagagagggagggagaggggggggttcgctagctctctctctcattggttaatgtttaactctgttgacatgttttcagagcg-----cgatctggtgatgct-tattgtacgttggagcttctgacacctatttttc
chimaera tbx3  ------------------------------ccccgttgcttcgccattggttctttcttaatgatgttgacatatccacacggc------taaagcggtgatgcttgcctgtacgttacagtttccgacacctttt----
lancelet       -------------------------------------gccctctcattggtccattcttaatgatgttgacattttcgcacgat------tgatacgataatgtc-ataagcacgttgtgggttccgacaccttt-----
acorn worm     ------------------------------cccatttggcgctgtattggtggatacttaattatgttgacatttcgccacgat------tgatgcgataatgtc-ataagtacgttgtgcgttccgacaccttgtt---
velvet worm    -----------------------------------------------------------agctgtgttgacatttcagcgagat------tg-cacgataatgtc-ataagtacatttggtcttgcgacacctctcctt-
abalone        --------------------------------------------------------------------------------------------------------------------------------------------
bootlace worm  --------------------------------------------------------------------------------------------------------------------------------------------

                                                                 GC-box_             CCAAT                    CCAAT
human TBX2     ----cccaggcccccctcccggccggagcttggcctgagctgtcaaaccccgcccccggagacccacaattggtccaaaaagc-gtaaaatcagcaatcaaggggggcctggctcgttagcgcaggggatccgagctg
human TBX3     ----tataactcagctctatcc-----cccagcactcgacctgtgaaaaccacgcctatgcagcaacacaattggtccgaaagc--gtcaaagagccaatcaagaggcctccggctccc---------------------
chimaera tbx2  ccctcttaactcatccat-----------cctggctctagttgtgaaaaccacgcctatacgggcacacaattggtccgaaagc--gtaaaatcgccaatcaactgtgc---ggctcattagcagcagagagccgagggg
chimaera tbx3  ----tataactcatctat---------ccctgggtcttagttgtcaaaaccacgcctatggaggcacacaattggtccg-aagc--gtcaaatagcctatcagtagggttctggtgctc---------------------
lancelet       ----tatagctcatatat-------------cttgtaaacttgtcaaacgcacgcctctctggcgaggggattggctgcaacgc--gtaaaatagccaatcaaggggacc--gtctcattagtg-----gctggcggcta
acorn worm     ----tacggctgatacgt--------------cgtgtaagttgccactaacacgcctatactcaacgctgattggtcgattcgt--gcgaaatagccaatcaactagcaa--acctcatta-------------------
velvet worm    ----tgtagagtagctat-------------cgtgtaaaactgcgaaaaacccgcctatttagcgcccggattggtgggaatca--tcgaaatagccaatcaacaagaac--ggctcattagcactagagttgggtgcga
abalone        ---------------------------------tgtgtacttgtcaaaaccccgccccgctcgttaaatgattggttgagaggcagatagcgcggccaatcggtggcgcc--ctttc-----------------------
bootlace worm  ----------------------------------gtaaacttgtcagatacacgcctatttctcaccctgattggcccgagcca--aagcagcagccaatcagattgctc--gtttcattaat-----------------

human TBX2     ggcagga-----
human TBX3     ------------
chimaera tbx2  ggaagca-----
chimaera tbx3  ------------
lancelet       gtcaccccgggt
acorn worm     ------------
velvet worm    ggcagcacgggt
abalone        ------------
bootlace worm  ------------
```

**Figure 15:** *Tbx2/3* promoter

```
                                                                    DUX________             E2F_________
human TBX3     tggccaatgtttaacccgaatgcagtggccaccaggccgcttttgtttgttcgcaaattaatcaccca--gcgcatggccggcgccagagtgggtttatcacccaccgggagggggcgcgccgggcacgcagagacaaa
chimaera tbx3  tggcgaatgtttaactaaaatg-gctcgccatcaattcccttttgtttgcttgcaaattaatcatccc--atacgcagttggcgccagtgtagatttatcacacacatgagggtgaatgagccgggcgaggagagacaaa
lancelet       gagaggaa---------------aatgacatacaattcccatatgcttcccccaaaattaatcatcca--atataccggtggcgccatggcgatttatca-catgtgcggtggtgaacgcgcgactgagagac-cacaaa
acorn worm     ggggtgga---------------aatcgtgttctattcccatatgttttccttgaatctaatcagaca--atgcgctactggcgccagggcgatttagc--gcgaatgtagcctgaataggcggcgtgctggcacacaaa
sea urchin     -------------------------------------------tgtttgtatagaaattaatcatccaccgcgcagccctggcgccagtcagctttagtcgtataaaagggaatgaataggtagccgtggagaatacaat
velvet worm    -------------------------------tttacctccttctgtttcactggact---atcagccgcgatctgttagtggcgccgtccagatttagtt-ctggtgcgggcgtgtagttatggtggtgccgc-tacaaa
shamisen shell --------------------------cgtgcagtattcccatctgtttttcagtagattaatcggcca--gtgtggggatggcgccacggcgatcgtgtt-caaacagtacgccgtatttgcgggtcgtgcctacacaat
bootlace worm  -------------------------------------------------------------------------------tggcgccagcggacttagca--gtcaacatagtcagtatttgtgcgtcgtctgt-cataag

                                                TBX_______
                                           PHOX_______
                    TBX_______        POU4____________
human TBX3     gaggagttcgcaccttcccgcttttgatcccagaattac------------------------------------ggcggcctccctgcctaatacgagcctcctggg--------------
chimaera tbx3  ga-gagttcgcaccttcctgcttttgatcttag-attag------------------------------------ggcaccctcccttcataatagaagtctccaggg--------------
lancelet       gg-cctttcgcacctcggctgtttgatctcgc--cttattaattatgttaacacttaggcgaatcctacagaaccaacaggctggctggccgcggccatctgattggtcgtgggtggggca
acorn worm     gg-ggcttcacacctcacgcgttttcactggcg-cttattaattatgttaacactttgcccagtc----------aacagcaattatccccaacgtatagccgattggtttagggtggggca
sea urchin     aa-gccttcacacctcacgtgttttctaccgcg-cttattaattatgttaacacttcggcaagtc----------aacaacaattatccgcgaaata-------------------------
velvet worm    ag-agattcacacctccgctgccttactcgcta-cttattaattatgttaacacttcacccagtc----------aacaacaattatttgacgaacaattgtgattggttcaaggtggagca
shamisen shell ac-cctttcacaccttgcctgtttacgggccta-ctttgtaattatgttaacacttgaatgaatc----------aacatcttttcttccgatcacaatactgattaaaccagggcggagta
bootlace worm  ga-gcattcacacctccactgttttatcttctg-cttattaattatgttaacactcgactgagtc----------aacaacaattat-----------------------------------
```

**Figure 16:** *Tbx3* enhancer

```
human          -tacattgctatttgta----aagtttgccttcacagcatttca-----gatgctgttggacttca-tgtccccaacct-------------agcttggtgaggg
chimaera       ------------------aaaagcgaggccttcgcagcacttta-----ggtgctgttggacttcactgtccccaac-----------------------------
chimaera       tcattttattcatttcc------aagggctttcacagctttttt-----actgctgttggatttctctgtccccaaactt-------gtagcaagtttggtgaagg
lancelet       ------------tttctgaagggtggaattttcacagcgctctc-----tgtgctgttggacctcagagtccccaacccctt----------ggggttcggcgaaaa
lancelet       -tacgttgct-gtttctgaagggtggaattttcacagcactctc-----tgtgctgttggacctcagcgtccccaacc----------------------------
lancelet       ------------------aagggtggaattttcacagcgctctc-----tgtgctgttggacctcagcgtccccaacc----------------------------
lancelet       ------------------aagggtggaattttcacagcgctctc-----tgtgctgttggacctcagcgtccccaac-----------------------------
acorn worm     tccttttctttactttc----aggtgtgctttcgcagcaatttt-----actgctgttggacttc---gtccccaaacc-------------acgtttggtgaaag
lobster        ------------------------------ttcacagcctgtcac----ggtgctgttggacctctcagtccccagccccctgaattttggggggcttggtgaa--
tick           ------------tctctcggggccgcccctttcacagcgccc-------atcgctgttggaccccc-cgtccccaacccc---ggcacagtggggttcggcgaaaa
cactus worm    -----------------------------tttcacagcgccccctagcgagtgctgttggacgagagtgtccccaacc----------------------------
```

**Figure 17:** *Znf207* 3′-UTR. This matches two nearby places in the chimaera genome downstream of *znf207*, and four nearby places in lancelet downstream of *Znf207*.

```
                                     miR-130                                                                                                              miR-92_
                                                                                                                                                               Let-7__
human CPEB2    ccattttgacatttgttatgcactatttttatatctctgtgagagatttttccaacagtcagc--tatttta-------------------tggcacactttttttgactgatgacatctc-ctttgctatacctcaa--
human CPEB3    -----------tttgttatgcactactactttttgtat-tctagacagttttcaaaagttggcagatttttttttttgtttttaaggaaaaaggcatgc--tttctgactgacatgatc---ttttgcaatacctcaa--
human CPEB4    --------atatttgttatgcactacttttgtatatct------tgtttttccaacagt-gaa--cattttt-------------------aggcacact-tttcactgacgggatatctctttatgcaatacctcaa--
chimaera cpeb2 ccatattggcatttgttgtgcactacttttgtatcttggtgagagttttttccaacagtcagc--tgtttta-------------------gggcacac--tttttgactgatggcatctc-ctttgcaatacctcaa--
chimaera cpeb3 --------acatttgttatgcactactttttgta-----tctggacggttttcaacagtcagctgatttttt-------------------aagcgtgc--tttctgactgacatgatc---ttttgcaatacctcaa--
chimaera cpeb4 --------acatttgttatgcactacttttgtatatct------tgtttttccaacagt-gga--tattttt-------------------cagatcact-tttttcacagatggcatctctttttgcaatacctcgatt
lancelet       -----------------------------------------------------------------acttttt-------------------aggcaggc--tttccaccta--tgtacc---atttgcaatacctcaagg
cactus worm    --------------------------------------------------------------------------------------------------------------------------------------------
shamisen shell --------------------------------------------------------------------------------------------------------------------------------------------

human CPEB2    -tttttggaatttagagaagaaatcagtagttttgc--aatgttaattatttagatattt--------------------------------------------------------------------------------
human CPEB3    --ttttgaaatataggaaagaaaatgatattttctaagtaagtttattc---------agaaaaatatatattaatttaattctgca-agaaaatgattta------acagatgggta------actttatttttttcat
human CPEB4    -tttttcatattgcaaagagtagc------tttttg--tacttttattactgagagatcttcatatac-------ttcattttttaa-tataaataatttt-----aataaattttat------tttcttatattctgct
chimaera cpeb2 -ttttctgaatttagaagagaaatcgatagttttgc--tatgtttattattgaaagatctgcatatata-attaatttaattcagca-aataaatgattta-----acaaaattgata------tttttatttttcatac
chimaera cpeb3 --ttttgaaatataggaaagaaattgatattttctaagtaagtttattc----------taaaaatatatattaatttaattctgca-aggaaatgattta------acagatgggta------attttatttttttcat
chimaera cpeb4 ttttttcatattctaaagagagatgaatagtttttc--tatgcttattgttcaaagatctgcaaatatg-gttaacttaattgtgca-aagaaatgagttg-----aagcaatcagtt------ttgctttctttcgttt
lancelet       ttttccagaattt---gagcaaatagatatttt-----gaagtttattcttgaaagttaggaatgagagtattaatttaattctgca-----gatgatttaaagagaagatgtgggta------attttatttttcttgt
cactus worm    ------------------------------------------------------------ggagtgacattattatttaattctgca-atcaaacagttgaaaaataaaatatgggtaagattgattttatttttctcgt
shamisen shell -------------------------------------------------------------aagatgtatgttaatttaattcagcacacaagataatttatggtaggaaaatgggta------attttatttttt----

human CPEB2    ----------------------------------------------------------------------------------------------------
human CPEB3    gct------------------tatttgtgtttt-----ttgaaaa-------tgaagaagttagagctttataactataatattaaagatcatatctaac
human CPEB4    ttt------------------tatacattt----------------------------------------------------------------------
chimaera cpeb2 atc------------------ttgctttct----------aaatt-------tggaagggtctaaacatttgta--------------------------
chimaera cpeb3 gct------------------tatttgcttttt-----tggaaat-------tgaaggagctagagctttataactttaatatggagaattgtagttagc
chimaera cpeb4 tat------------------tattttttt----------------------------------------------------------------------
lancelet       att------------------gttctgctttttgcatatgaaaac-------tggaaagctttaaacttttttattata---------------------
cactus worm    aatattattatgtatgattccttttcagcctttgcctgtaaggatgccagtgcaggagattttaaacttttttattata---------------------
shamisen shell ----------------------------------------------------------------------------------------------------
```

**Figure 18:** *Cpeb2/3/4* 3′-UTR. Previous studies showed regulation by miR-92 [11, 12] and miR-130 [13], and predicted these Let-7 and miR-92 binding sites [11].

```
                                                                                                                                                                          PKNOX_
human ID1           --------------------------------------------------------------------------------------------------------------------------------------------
human ID2           ---------------------------------------------------------------------------------------------------------------------------ggcgcgcgcgctgagtg
human ID4           -------------------------------------------------------------------attgtgc------------------------------------------------------------gggctg
chimaera id1        -----------------------------------------aaggtgacaa-attaaagggcgcggtatcagggt-----gcagatgattgccagttaattgctgccccgcg-cggcgcccttgtagtgcg-gctgattg
chimaera id2        ---------------------------------------------tgatgg-------------ccgagcagcc---------gctgccc-----------------------cggtgcggccggcggggccctcaactg
chimaera id3        --------------------------------------------------------------------------------------------------------------------------------cgcttttgatgg
chimaera id4        --------------------------------------------------------------------------------------------------------gggggggg-ggccgggttggattgggg---tgattg
lancelet            ----------caaatgc-acaggtgcgcttggaagggggcgaaggtgacaacattagggctccatttatcaaca------aggatcggc-------------cgcaagtgta-cgggcgccttgtgcgcggatttgattg
acorn worm          ---------aggtgtgt-aagtgtgaaatgacaacagttcgaaggtgacga-attaaaaagccatttatcacaca-----ggaattggtcccgagtaattgccagtgc--cg-cggttcccttgtactaagttttgattg
sea urchin emc      -----------------------------------------aaagtgacga-tcccaaaagccatttatcgctgctctcgatagtttgttagtaattgat--tcaggctgcg-ggacgcctttgatgttgg---tgattg
sea urchin emc-like -----------------------------------------aaagtgacga-tcccaaaagccatttatcgctgctctcgatagtttgttagtaattgat--tcaggctgcg-ggacgcctttgatgttgg---tgattg
velvet worm         ---------aggtgtacagggtgtgaggtatcactaggccgaaggtgacga-attctaatacaagtcagcagtc-----------------------------gtcagtacg-cggtatatttgaagtataccctgattg
cactus worm         -------------------agcgagaagtaccaccaggccggaggtgccgg-gtgggatcgccgtttatcgttc-----------------------------tgcgatgtt-cggaacatttgatctcggccgtgattg
chiton              ---------aggtgtgc-gggtgtgaagtatcaacgggccgaaggtgacga-tttttaatgcaatttatcagcc-----------------------------tctcgcggg-cggcacctttgatgtaaa-catgattg
sea hare            ---------------------------------------------------------------------------------------------------------------g-cgggccctttgaagtggtggctgattg
abalone             ---------aggtgtgc-ggctgtgaaccatctacagggcgaaggtgacgg-attttaatgcaatttatcagcc-----------------------------atgggaggg-cggcctctttgatgtttgtgttgattg
owl limpet          ---------aggtgtgc-aagtgaggagtaaccaagggccgaaggtgacga-attttaatgcaatttatcagtg-----------------------------agtcgaggg-cggcttctttgaagttatgtttgattg
oyster              caaatacgcaggtgtgc-tagtgtgaagtatcagggggccgaaggtgacga-attttaatgcaatttatcagta-----------------------------acttcaggg-cggcagctttgaagttgtgtgtgattg
nautilus            ---------aggtgtgc-tgcggtgaaatatctgcgggccgaaggtgacaa-atcttaacgcaatttatcagtg-----------------------------aaggaagcg-cggcttctttgatggtgcgcgtgattg
peanut worm         ---------aggtgtgc-gagtgtgaagtatcaacaaactcgaggtgtcgc-ttgtgaatgcaatttatcagtc-----------------------------acctttggg-cggcccttttgaagtcacccttgattg
sandworm            ---------------------------------------------------------------------------------------------------------------g-cggcgcttttgaagtgcacagtgattg
Satsuma tubeworm    caaatttgcaggtgcgctgcgaccgaggtatcggctggccgaaggtgacga-attcgaaggcaatttatcagcg-----------------------------gacttaatg-ctgccgctttgatgtaggcgttgattg
shingle tubeworm    ---------aggtgtgc-atgtgctaagtatcaacaagccgaaggtgacga-attctaataccgtttatcagta-----------------------------agtaaagag-cggtgtatttgaagtaagtcatgattg
shamisen shell      -------------------agtgtgaaagatctacaggccgaaggtgacga-attgtaatgcaatttatcagtc-----------------------------accggtgag-cggaggctttgatgtacaccttgattg
horseshoe worm      ------------tgtgt-atgagtatgctgacactaggccgaaggtgacaa-aacataatgcaatatatcggcc-----------------------------taccccgtg-cggttcttttgaacttagtgttgattg
bootlace worm       ---------aggtgtgc-tagtgtgaagtatcatgaggccgaaggtgacaa-attctaatgcaatttatcagtc-----------------------------acttgagggtcggttccattgaagtttactatgattg

                                                                                                      Atf_____
                    ____      RFX______________                                         ZNF24__________             Plagl1___
human ID1           --------gtcgtctccatg-gcgacc---gcccgcgcggcgccagcc-tgacagtccgtc-cgggtttt-atgaat-gggtgacgtcacaggcc-tggcgtctaacggtctgagccg----------------------
human ID2           acgg---cgcggttgccatg-gcagcc---gcctgagcggcgccgcga-ggacaaggctgc-agggcggc-gtgaat-gggcggcgtcacgcgcc-tggcgccagagagtctgctccggg---gctccggctccggcccc
human ID4           gcgt---ctgcgttgtcagg-gcgacc---gcccggccggcgccagcg-gagcgggggagg-cgggctcc-gtgaat-gggtgacgtcacgcagc-cggcgccaggcagtctgggcgg----------------------
chimaera id1        gcgg---gagcgtcgccatg-gagacc---gcctgcccggcgccagcg-tgacagc-cgtc-aaggccgc-atgaat-gggtgacgtcacgggcc-tggcgcctgccagtctgagg---------gctgcggggacctgg
chimaera id2        acagc-cgcccgttgccatg-acagcc---gcctcggcggcgccgcgc-ggactggctcga-gggccgct-gtgaatggggtgacgtcacggggc-tggcgccggcgagtcggcgcgg-------ggccgaggggacccg
chimaera id3        gcagccccggcgtcgccctg-gaaacccccgccgccgcggcgccaggg-c------------tggcgctc-atgaat-gggtgacgtcacggggc-tggcgccaggctgtctggagcggggtcgggctgcg---------
chimaera id4        gcgtgcgacgcgttgccacg-gcagcc---gccgggccggcgccagct-gga-----------gcgctgt-gtgaat-gggtgacgtcacggggc-tggcgccggccagtctgggcgc-------gcgctctgttacgat
lancelet            gcgt---ggtcgtcgccata-gcgacc---gccagcccgacgccagtt-tgacaggcatca-agcg-ggg-atgaat-gggtgacgtcacgacac-tggcgccagagaggcgcgcctg-------gctgttgttgattgc
acorn worm          gcgt---gtgcgtgtccata-gcgacc---gccatcgtgtcgccagtc-tgacagcattca-tgagccca-atgaat-gggtgacgtcacaagac-tggcgccagcgagccggcggca-------gctgttgttgattgt
sea urchin emc      gcgt---gtgcgttactatg-gcgacc---gccagaccggcgcccgtc-tgctatcggatt-gtgggtgg-atgaat-gggtgacgtcacagatcgtggcgccggtctgtcgccgtgagc-tctggtagtattggagggg
sea urchin emc-like gcgt---gtgcgttactatg-gcgacc---gccagaccggcgcccgtc-tgctatcggatt-gtgggtgg-atgaat-gggtgacgtcacagatcgtggcgccggtctgtcgccgtgagc-tctggtagtattggagggg
velvet worm         gcag---cttcgttacgctg-gcaacc---gccagtctggcgccactg-ggatggcttaca-ctctttga-atgaat-ggatgacgtcacatagc-tggcgccactcagacggtggca-------gctgttgttgttcgt
cactus worm         acag---gcttgttgtcttg-acgacc---gccagcctggcgccagac-gaccagcccggc-cgtcttgt-atgaat-aggtgacgtcacaggtc-tggcgccagcctggcggagacg-------ggtgttacagattgt
chiton              acat---gtgggtaacgtta-gcgacc---gcctggctggcgccgagt-ct-----------caagcctc-atgaat-gggtgacgtcactcgtc-tggcgccgctcagtcggctggg--------ctgttgttgattgt
sea hare            acag---tcgtccagtggcgtgcgacc---gcctgcctggcgccagta-acaagcaaaaac-tcggctccactgtat-gcgtgacgtcaccatgc-cggcgccagtctgtctggcccg-------actgtt--tagaccg
abalone             acag---cgtggtagctaca-gcgacc---gcctggctggcgccgtctacgatagcaaata-tacacaca-gtgaat-gggtgacgtcacagagc-tggcgccagtctgtcgcgcttc-------gccgccgctgattgt
owl limpet          acag---gtatgtaacgcta-gcgacc---gactatacggcgccggct-cgagaggaaata-catacagtgttgaat-gggtgacgtcacaagtg-cggcgccagtcagtctgctagc-------tgaattgttgattgt
oyster              gcac---gtcaaaaacgcta-gcgacc---gcctgagcggcgcctaga-ccatagcaaagagtatgaaag-aagaat-gagtgacgtcacactgt-cggcgcctctctgtc-----------------------------
nautilus            acag---gccggtaacgttg-gcgacc--agtctgcatggcgcccgcg-cgatagcgaaca-cacactac-atgaat-gggtgacgtcacggggc-tggcgccggtctggcgcggaga-------accagtgttgattgt
peanut worm         acag---tagtgttgccttg-gtgacc---gccatactggcgccgagt-caatggtatata-catatact-ttgaat-aggtgacgttacatgga-tggcgccagtgagtcgcgctca-------gctgtgg--------
sandworm            acag---tgcttcaggtggg-gtggcc---gccagcccggcgccagtt-cgccagccaaga-catacacc-atgaat-gggtgacgtcacatagc-cggcgccgctgcgtcgccccta-------cctgctcacgattgt
Satsuma tubeworm    acag---acgcgtcgtcatg-acgacc---gcctgcctggcgccagtt-cgatggcccta--tattgcgt-ctgaat-gggtgacgtcacgatgc--ggcgccagtacggcgccgctg--------------------tt
shingle tubeworm    acag---gcttgttgctata-tcaacc---gcctggccggcgccggct-catagtg---------------tagaat-ggatgacgtcacgaaac-tggcgccgtcctggcgggacca-------gctgaga-ggaaaag
shamisen shell      gcag---tgttgttactata-gcgacc---gccggcccggcgccgagt-cgatggc-tgta-tacacagt-atgaat-gggtgacgtcacacggc-tggcgccacactgccgggggca-------gctgttgttgattgt
horseshoe worm      acag---gtcggtgaccata-gctacc---gccagcctggcgcctaga-ctatacc--tta-tatactgg-ttgaat-aggtgacgtcacagcac-tggcgccggccagctgggcctg-----------------attgt
bootlace worm       acaa---gagtgcttggtta-gtgacc---gccgtggcggcgccaagt-cgatagtctgta-cacacact-atgaat-aggtgacgtcacgagac-tggcgccggcccgtcgcagtcg-------gtcggacagaactat

human ID1           ------------------------------------------------------------------------------------------
human ID2           gccgcggcc---------------------------------------------------------------------------------
human ID4           -------ccgagccaatgaatcagctgtccggggg-------------------------------------------------------
chimaera id1        ggggcgct---------------------------------------------------------------------------------g
chimaera id2        ctcggctccg--------------------------------------------------------------------------------
chimaera id3        cgctgattgctgccaatacatcacctgtca------------------------------------------------------------
chimaera id4        cccgc-----tgccaatgaatcagttgtcagtgag-------------------------------------------------------
lancelet            gccgtgttcgcccaaattaatcacatgtcaaatatgattgtg-gctgtcatgcgaaa------agtacacacagcccgggtgcagctgtg
acorn worm          ggcttgttttccccaattaatcacttgtcaaatctgattcgg-tatgtcatgcgaaa------aatacacacacctctcacacagctgtg
sea urchin emc      ggcttaggagggccaattaatcacctgtca------------------------------------------------------------
sea urchin emc-like ggcttaggagggccaattaatcacctgtca------------------------------------------------------------
velvet worm         ggcttgttctccccaattaatcacttgtcaaaccagattggc-cctgtcatgtgaaa------aacacacagagcacggat---------
cactus worm         agcgtcttcagtacaattaatcacttgtca------------------------------------------------------------
chiton              gacttgttctcgccgattaatcacttgtcaaagcagacttgcggagctcctacgaaaag-------------------------------
sea hare            ggctccccaagcgcaatttaccacgtgtcaaagccgatttag-cacgctgcgtgaaa------acctcactaaacacaggtgc------a
abalone             agagcgt--agagctattaatcacctgccacctctcgccgcg-actgacaaggggga---------------------------------
owl limpet          ag-aggttcacatcgagttatcacgtgcagaaaacgatagaa-aatgtcatgagaaa------aat------------------------
oyster              ------------------------------------------------------------------------------------------
nautilus            tagttgtactccccaattaatcacttgtcaaatcagattctg-aatgtcatgtgaaagtacacaatccgcagagtctgagtgc-------
peanut worm         gccagctcttgtcactctaatgacttgtcaaagcggatt---------------------------------------------------
sandworm            -cccccttcgcgccaattaatcacttgtcaaatcagattgaa-ggtgtcatgcgaaa------agtacacaaac----------------
Satsuma tubeworm    ggatttactgacggaattaatcacttgtcaaatcagatcggtcggtgtcaggcgaaa------cacacgc--------------------
shingle tubeworm    gggtcgttctccataattaatcacttgccaaattt----------tgccatgagaaa------aatacacaaattt--------------
shamisen shell      tgcttgttctcgccaatttatcacttgtcaaacctgatcggg---tgtcatgtgaaa------aatatatacac----------------
horseshoe worm      agcttgtaccccattattaatcatatgttaaagaagat----------------------------------------------------
bootlace worm       ggcatgttcttggctattaatcacttgtcaactatgcttata-cttgtcccatgaaa------ag-------------------------
```

**Figure 19:** *Id* enhancer

```
                                                                                            E2F___________
                              MEIS____               CCAAT                               HOX_______           HOX_______     E-box_
human LIN28B         --------------------------------------------------cccttaa-ggatacgaggtgaaattagtagcaagaaagcatgtaattgacaaagtcacgtgtgctcagggggccagaaactggagagagg
chimaera lin28b      ttatgtgcggtgacaggttggagcagagtgggattggctcggcgctgcctctccgaa-ggctgcgatgtgtaattagcggcaagaaagcatgtaattgacggagtcacgtgcgccag---ggccaggaa----aggagga
chimaera lin28b-like ttatgtgcggtgacaggttggagcagagtgggattggctcggcgctgcctctccgaa-ggctgcgatgtgtaattagcggcaagaaagcatgtaattgacggagtcacgtgcgccag-----------------------
lancelet             ttattctcaatgacaggtagcagagccacgctattggctgttgccagggcggccgct-gattggtatgtgtaattagtggcaaagagccatgtaattg--tacgtcacgtgtagcag-----------------------
velvet worm          --------------------------------attggttgttgctaagataataattccactagtatttgtaattagtgccaaagaagcatgtaattg--tacgtcacgtg-----------------------------

human LIN28B         agagaaaaaaatcaaaagaaggaaagcacattag
chimaera lin28b      aaaaaaaaggggctaaagaggaaaagcacgttag
chimaera lin28b-like ----------------------------------
lancelet             ----------------------------------
velvet worm          ----------------------------------
```

**Figure 20:** *Lin28b* promoter

```
                                                                                                          MRE_____
human MEX3B      ------------tgctttctaac-----------atttgtataaatag-aat--tcagtgcaagttacaaaaattctgttgcaccac-tctagtttt-agtatttcta-ttttaata--catttgtttaccacttgttta
human MEX3C      ----------------------------------------------------------------------------------atcaa-tatagttta-gtatttgctaattttacta---cactct-------tttgtta
human MEX3D      ---------------------------------------------------------------------------------------------ttta-attttttttt-ttttttac---ttttacttttta-cctcttg
chimaera mex3b   aaggaacagttaagctttctaac-----------atttgtacaaatagaaaa--ccagtggaaattac--aaattctgttgcaccac-tatagttta--gtatttcta-ttttaata--catttgtttaccacttgttta
chimaera mex3c   ccg----------------------------------------------------------------------------------------------------------------------------------tttgtta
chimaera mex3d   --------------------------------------------------------------------------------gcaccag-tacagattactaaatattca-agtttatc---atttgtttacct-cttactg
lancelet         ------------tgctttctaac-----------atttgtataaaata-gct--acacatgccgatacgataatttcgttgcac----catagtctaggcatgtacta-ttttaata---tattatttacaa-cagttta
acorn worm       ----------------ttctaacatat-tatattatttgtataaatag------caactgactatatggtaaaatccgttgcact---catagctag-aatttacttatttttaata---tattattta----cagttta
velvet worm      ----------------------------------------------------------------------------------------ataggttta-gtattttcta-ttttaata---cattatttacca-cagttac
cactus worm      ----------------------------------atttgtata-----------gtagcgaccgacttgtatgatttgttgcacat--gttagttta--gtatttcta-ttttaata---tattatttacta-ccgtttg
chiton           ----------------------------------atttgtataaatag-ca----tttcaaaatgaccatttttgttgtttcacact-catagttta-gtattttcta-ttttaatatgatattatttaataccagccga
abalone          ---gaagatttaagctttctaacatacatatattatttgtataaatag----------attctcaaagacttaatttgttgcacatc-catagttta-gtattttcta-ttttaata-----ttatttacta-ctctgtg
owl limpet       -----------------------------------------------------------------------------gttgcactta-aatagttta-gatttttcta-ttttaata--ttattatttactatttgactg
nautilus         aaagcaagattatgctttctaac------attatatttgtataaatag-------------tatctgaaaaaaatttgttgcacaac-catagacta-gtattttcta-ttttaata-----ttatttacgaccagtctg
octopus          --------------------------------------------------------------------------tttgttgcacaacttagagacta-gtattttcta-ttttaata------tatttacgagctatctg
Satsuma tubeworm --------------------------------------------------------------------------------------------------------------------------------------------
shamisen shell   ---------ttatgctttctaatg----catattatttgtatgaatag-------caaaactgacaacaattaatttgttgcacaac-catagttta-gtattttcta-ttttaata--ttattatttatcacaattctg
pale anemone     a--------------------------------tatttgtacaagata-tgtagtcataatttggatacgatattccgttgcactac-tatagttgg-------acta-ttttaata---t-------------------

                                                                 MRDE__________________________________
human MEX3B      tgtatatgtaggtgatgttacttgagcttaaatgtacttta--------ctgagcaaagtttaaaaaaca---aagta-----tattttatttt----atgataaag-ggcctttaacct--catggtcaaat--actaa
human MEX3C      tgtatatgtagggaagtcatagggattataaattcaattt-----------gagtaaaatttaaaa-----------------ccatatatttt----atgataaag-ggcctttaacttaagatggccaaag-cactga
human MEX3D      tgtatatgtagggaatttata---gggaaatatgtacttta--------tggaataaattttaagaacta---aaata-----tattttattttaaataaagtaatg-gacctttaatcttacacagctaaat-tactga
chimaera mex3b   tgtatatgtaggtaaagttacttgagcgtaaatgtacttta--------ttgagcaaagttt-aaaaacg---aaata-----tattttatttt----atgataaag-ggcctttaacct--catggtcaaat--actaa
chimaera mex3c   tgtatgtgtagggaagtgataaggattataaattcaacctattgagtcgaaacaaaaagtttaaaa-----------------ccatatatttt----atgataaaatggcctttaacttaacgtggccaaag-cactga
chimaera mex3d   tgtatatgtagggaatttata-ggactgaaaatgtacttta-------tttgaataaattttaaaaaaaac-taaata-----tattttatttt----aagataatg-gacctttaaccttacatggctaaag-tactga
lancelet         tgtatatctaggtgtgaaaca-tgaccgcaaatatactatg-------gagaaaaagtttactaaataca---aaata-----ttttttatttt----gggcctggt-aaccatg-------tttggccagatatactaa
acorn worm       tgtatatgtaggtgtgagata-tgaccaaaaatg---------------ttgaacaaagtttaatacaaatgcaaatat----ttttttattttttgtttggccagct--------tactgaaatggccagatatactaa
velvet worm      tgtgtatattagtatagaact-ttatcaccagca---------------aaaaaatgaatactaaaga---------------tattttatttt-ataaggtccatt-----------tcgttgtggccat---tactaa
cactus worm      tgtatatgtaggtgtga---------------------------------------------------------------------------------------------------------------------------
chiton           tgtgtatatgtagagaaactc---aatacagatgttgtaaa--------caagtgaaaagtctaaaaaaa---agata-----ttttttatttt---caaggccaat-gcaaatg----t--tgtggccagat--actaa
abalone          tatat-tgtagatgttccaca--gattaccaataactcgtc-----------gacaaagtacaagaa------agata------tttttatttt----gaggccaag-caaaata---------tggccgtt---actaa
owl limpet       tatatgattgtagacgtttat----------------------------aaagacaaaataccatataag---atgaaaagatatttttattttaataaagaccact-aaataaa---------tggtcgat---actaa
nautilus         tgtatatgtaga---------------------------------------------------------------------------------------------------------------------------acaag
octopus          tgtatatgtaga---------------------------------------------------------------------------------------------------------------------------acaag
Satsuma tubeworm -------------------------------------------------------------------------------------ttttatttt----atggcaata-tgcagtg---------tggccaaa---actaa
shamisen shell   tgtatgtaggtgtgtgactatggaacccaaagtaaagatta--------agatgcagagtttaaagttaa---atacaaaga-tattttatttt-gttaggctcagc-taataag----------agcctgta--actaa
pale anemone     --------------------------------------------------------------------------------------------------------------------------------------------

                    miR-374
human MEX3B      ---tattatatttgctgagaca---agatttgaaa------------ttgtatcaagagttttatttttctgacattta---------------aagttctacataataaaggtaaaact
human MEX3C      ta-ttatatatttgctgtaaag------------------------agaattataagagttttatttttctgatattaa-aagttacttaataaagacttgtttccattaacttgaa---
human MEX3D      ---ttatatatttgctgagctgatttaagggttaa-------aaaaattgtatcaagagttttattttttgacttcaaa--gccttcttaataaagcctcttttctacatgt--------
chimaera mex3b   ---tattatatttgctgagaca---agaattgaaa------------ttgtatcaagagttttatttttctgacattta---------------aagttctacataataaaggtaaaact
chimaera mex3c   ta-ttatatatttgctgtaaag------------------------agaattataagagttttatttttctgatattaaaaagttacttaataaagac-cgtttccattaaagtgaa---
chimaera mex3d   ta----tatatttgctgaactgacttgatggtatg---------aaattgtattaagagttttattttttgacttaaaaaaaccttcttaataaagactttttacagtatgt--------
lancelet         tattattatatttgttgagact--tgaggattata------------aaattgtaagagttttatttttctgccattaa---------------agttgtattacaataaatgacaagct
acorn worm       ---tattatatttgccgagatt---tgagattatt----atggaattgttattaaaaagttttatttttctgtcattaa---------------agtaaaagttct--------------
velvet worm      ---tattatataaactaagaga-----------------------tatacaatttaaagttttatttttctgccagta------------------------------------------
cactus worm      ------------------------------------------------------------------------------------------------------------------------
chiton           ---tattatatttgctgaactc---aaatattata-------------aatgtaaaaggttttatttttctggcattaa---------------aa------------------------
abalone          ---tattatatttgctgaaact---cagaagtata-------------agatgtaagagttttatttttctgccattaa---------------agtcaagttccaa-------------
owl limpet       tattattatatttgccggaact---caattataag-------------agatgtaaga-ttttatttttctgccattaa---------------agtaaa--------------------
nautilus         t-----------------------------------------------------------------------------------------------------------------------
octopus          t-----------------------------------------------------------------------------------------------------------------------
Satsuma tubeworm tgttattatatttgctggaccc---agaattttaa-------------gattgtaagagttttatttttctgccattaa---------------aaat----------------------
shamisen shell   ---tattatatttgtggagagt---aagaaaaaaagtataaaaaaaagaattgtaagagttttatttttctggcattaata-----------aaagttgtatttcaa-------------
pale anemone     ------------------------------------------------------------------------------------------------------------------------
```

**Figure 21:** *Mex3* 3′-UTR. MRDE: previously predicted Mex3b-responsive destabilizing element [14]. MRE: resembles a MEX-3–recognition element [15]. miR-374: previously predicted miRNA binding site [16].

```
                                                           GLI___________                  RFX______________                                             CC
human PTCH1    gaggctgccc-----cgaagccgaggatgcacacactgggttgcctacctgggtggtctc-tctac-tttggtgagctgttgctaagcagctaataatag---tcatgtttgctgagtaatttctgccct-ccgcgagcc
chimaera ptch1 --------------------------------tggatgcactcacagcctgggtggtctc-tctac-tttggtgagctgttgctaagcagctaataatag---tcgtgtttgctgagtaatttctgcct--tcctcggcc
chimaera ptch2 --------------------cccacccccccattgatgctctggcagtctgggtggtcctggccac-tttggtgagctgttgcttagcagctaataatagt--agatgtttgctaagtaatttgcttcttccctttagcc
lancelet       gaaggcgcctggcggtggaggatggatgtgtcagggtacccctacgacttgtgtggtctg-tcttc-tttggcgagccgtttcctagcagcaaatgacggg--ccgtgttactcgagcaaattgtggag--cccgcggcc
acorn worm     ----------------------tcctctttttatcctatcccaacttcttgtgtggtcta-cctctttttctggtgacgttactaagcaactaatgagaac--tcgtgttacatgagtcatttggccg---tgagtgacc
scorpion       ---------------------------cctctttggtggctaggcaacgtgtgtggtccggacgac-gtcggtgatccgttgcctagcaactaat---------------------------------------------
horseshoe crab -----------------------------gccacagcgtccccgcgacgtgtgtggtcttctccac-tttagcgaaccgttgcttggcaactaatgcaaggttcgctgttacgagaggagctagaaag---ccacgagcc
horseshoe crab --------------------cccgcattctcaacagcgtccccgcgatgtgtgtggtcttctatac-tttagtaagccgttgcttggcaactaatacaaggctcgctattacgagaggagctagaaag---ccacgagcc
chiton         -------------------------ttccctcttcgacctaccgcaatctgggtggtctt-tctac-tttggtgagccgttgcctggcaactaatgtgcagc-tgatgttacttgagcaatctgcgcta--taattgacc
sea hare       -----------------------------------------tcacttcttgggtggtcta-tctac-ttgggtgagctgttgctaagcaactaatgttgggc-tggtgttacacgagcaatcctactga--taattgacc
abalone        --------------------------------tgggttcccttccttcttgggtggtctg--ctac-ctgggggagctgttgccaggcaactaatgttgggc-tggtgttacacgagcaatcctactga--taattgacc
oyster         --------------------cttatcgcactgttggcggtcccgcaacttgggtggccgagcttctatgtggtgaacggttgcttagcaactaatg--------------------------------------------
shamisen shell -----------------------------ccgtttgttcctggccagcttgggtggtctt-tctac-tttggtgagctgttgctaggcaactaatgtcgggc-tgcagttacatgagctgcctgtgggc--ttaagaacc

               AAT
human PTCH1    aatcgcgtcgcagaaacccaagccatctgaca------------gccccgggggcggga----g
chimaera ptch1 aatgagatcccagcaatacaagccttctcccagccatctgacg-gccctggaggcgggtttt-g
chimaera ptch2 aatgggaatccggtcgagaggactttccatcaactctatctgatgggctggaggcgggtt----
lancelet       aatcagagcgcgggaaggtgacccctcgatcacgggagctcga-ggcggagaggcgggttttgt
acorn worm     aatcagattgaagaaaattgacccttctgtggggcc-gtttga-caggagggggcgtggtt---
scorpion       ----------------------------------------------------------------
horseshoe crab aatcgaagcg------------------------------------------------------
horseshoe crab aatcgcagcg------------------------------------------------------
chiton         aatcagagcgctgatctgtttcccatcaatcatg------------------------------
sea hare       aatcag----------------------------------------------------------
abalone        aatcag----------------------------------------------------------
oyster         ----------------------------------------------------------------
shamisen shell aatcatcgccgcgctctgcgcgccatcaatcacgtc-accgca-ttcccgggggcgggccttgt
```

**Figure 22:** *Ptch* promoter

```
                                                                                                                     RFX___________ NRRE_____
human              --------------aagggggcggagctttggcg------------------tggggcggccaatagt------------gggggt--ggctgcgtgggtcgccatggggacgggg---ctgttcccggggaggctgtg-
chimaera           --------------aacggggcggcgccggctgc------------------ggcgcgggccaatgag------------ggcggc--ccgtcgtttggtcgccatggtgacgggg---ttgttcccggga----cctg-
lancelet           ----gtgggttttgaaaattatagcttcaagtatgggctgtctcccgccaaacaaatggaccaatcagcgaagaccttttgacaag--tcatcatcgtgtttccatggttaccggg---tcgggctgggg-----cttg-
acorn worm         tcacgtgatgcaatgctaagttagtttctagatc--tccataccaattactgcgaactagccaatcaggatacactcttcgacgggtcatcaactgcagtttctatggtaacgggg---tcggactcggc-----cttg-
sea urchin         ----------------------agtttctagaat--ccctgaattaat-cagtgcttcgaccaatcaacgtcgcgtactgggcagt--tcacgatgtagtcgtcatagctacgggg---tcagtctctgc-----cttg-
centipede          --------------------------------tc--cccttagcaacc-acaccgaacgaccaatgagcgtgcgggctttgacagg---tccattgacgtttctatggtaaccggg---tcagtctcaac-----cttg-
millipede          -----------------------------------------------------------------------------------------caccacaatgtttctatagtaacggagtttttggtgtcgtc-----cttg-
scorpion           ----------------------------------------------------------------------------------------atcaattaatgtttctatggtaacaggg---tcggtgtcgtc-----ctga-
scorpion           ------------------------------------------------------------ccaatcacatagccgcatttgacaga-tcccaattgatgttactatggcaacaggg---tcggtgtcgtc-----ctga-
tick               ------------------------------------------------------------ccaatcacaaggcagtctttgacag---atcaattgatgtttctatggtaacatgg---tcggtgtcgtc-----ctga-
horseshoe crab     ------------------------------------------------------------ccaatcacgttactgcctctgacag---atcaattgacgtttctatggtaacaagg---tcggtatcgtc-----gtat-
horseshoe crab     ----------------------------------------------------------------------------------------atgaactgatgtttctatggtaacaagg---acaaaatcatc-----gtga-
horseshoe crab     ----------------------------------------------------------------------------------------atgaactaatgtttctatggtaacaagg---agaaaatcatc-----gtga-
horseshoe crab     ------------------------------------------------------------ccaatcacacagctgcttcccacag---atcaattgatgtttctatggtaacgagg---tcggtgtcgtc-----gtgt-
velvet worm        -------------------------------------------------------atgggccaatcacatttgagcttttgacag---gtcgattgatgttgtcatggaaacaggg---tcggtgtcgtc-----cttg-
chiton             ---------------------------------c--cccacaactactataattaaacatccaatcgcaatgtgccatttgacagg--tctacttgacgttgtcacggtaacgagg---tcagtgtcgtc-----cttg-
sea hare           ------------------------------------------------------------ccaatcaacgcgaagcgactgacagc--cgcagttgatgttggcaagacaacgggg---tcagtgtcgtc-----cttg-
abalone            ------------------------------------------------------aactatccaatagagacttgccatttgacagg--tctacttgatgttggcaaggcaacgggg---tcagtgtcgtc-----cttg-
owl limpet         ------------------------------------------------------aaccgaccaatcgagactagctatttgacagg--tctacttgatgttggcaaggcaacgggg---tcagtgtcgtc-----cctg-
oyster             ------------------agttggtttctagatc--gcagattctgctataattaacgaactaatcgtagaacgcctttcttcagt--tctacttgtggttgctatagtgacgggg---tcagtgtcgtc-----cttg-
nautilus           -cacatgatgttagaccggaccgttttaaggcac--tccaaacttactataattaactacccaatcgtaagttgcattttgacagg--tctacttgatgttggcaaggcaacgggg---tcagtgtcgtc-----cttg-
octopus            ------------------------------------------------------aactacccaatcgtaacttgccttttgacagc--tctccttgacgttggcaaggcaacgggg---tcagtgtcgtc-----cttg-
peanut worm        ----------------------------------------------------------------------------------------acccagtgacgtacccatggaaacgagg---tcagtgtcgtc-----cttg-
sandworm           -------------------------------------------cctactcagcggctcagccaatcggagcgcttctcatgacag---ttctcttgatgtcgtcatggtgacgagg---tcagtgtcgtc-----cttg-
Satsuma tubeworm   tcatgtgatattttgactcatgagtttctagtctc-cccacatctactaaaattactccaccaatcgtattgctccgtttgacaag-tctcattagatgtcgccatggtgatgagg---tcagtgtcatc-----cttg-
shingle tubeworm   -----------------------------------------------------------------------------ttcgacagg--tctttctgacgtcgctatggaaacaagg---tcagtgtcgtc-----cgtg-
shamisen shell     ----gtgatgtttgaacacgacagtttcaaggct--caccaatcctccgtaataactcgcccaatccaagattgctatttgacagg--tccttgtgacgtctctatgacgacgagg---tcagtgtcgtc-----cttga
horseshoe worm     ----------------------------------------------------------------------------------catg--tggtgatgacgtcactatagcaacaagg---tcagtgtcatc-----ctta-
bootlace worm      ---------------------------------------------------------tacccaatcgaaaggcgctatttgacagg--tcttcttgacgtcaccatagagacgggg---tcagtgtcgtc---ctcttg-

                                   PKNOX_______
                           MEIS____
                            TBX______________
                               E-box_
human              --at-gggttgacaggtg-cgtgacagt-------------
chimaera           --at-ggcttgacaggtg-cgtgacaatccagctcttca--
lancelet           --at-gggttgacaggtg-cgtgacagtccagctatccagc
acorn worm         --at-ggcttgacaggtg-cgtgacagttcagctacccagc
sea urchin         --at-tgcttgacaggtg-cgtgtcagtatagctacccagc
centipede          --atgggcttgacaggttgtctgacatttc-----------
millipede          --at-ggcttgacaggtg-cgtgacagttc-----------
scorpion           --at-gtcttgacaggtg-cgtgacagtttaacttgacagc
scorpion           --at-gtcttgacaggtg-cgtgacagtttaacttcacagc
tick               --at-gtcttgacaggtg-tgtgactgttcaaatccacag-
horseshoe crab     --at-gtcttgacaggtg-cgtgacagttcagcctcacagc
horseshoe crab     --at-gtcttgacaggtg-tgtgacagtttaac--------
horseshoe crab     --at-gtcttgacaggtg-tgtgacagttcaac--------
horseshoe crab     --at-gtcttgtcaggtg-cgtgacacttcggccacacag-
velvet worm        --at-ggcttgacaggtg-cgtgacagttcagctccacagc
chiton             --at-ggcttgacaggtg-catgacagttcagct-------
sea hare           --at-ggcttgacaggtg-cgtgacagttcagctct-----
abalone            --at-ggcttgacaggtg-cgtgacagttcagctctc----
owl limpet         --at-gtcttgacaggtg-cctgcgagt-------------
oyster             --at-atgttgacaggtg-cgtgacagttcagctccc----
nautilus           --at-ggcttgacaggtg-cgtgacagttcagct-------
octopus            --at-ggcttgacaggtgccgtgacacttcagctac-----
peanut worm        --at-ggcttgacaggtg-cgtgacaatccaact-------
sandworm           --at-ggcttgacaggtg-cgtgacagtttcagctc-----
Satsuma tubeworm   --at-ggcttgacaggtg-cgtgacagttcagc--------
shingle tubeworm   --aa-gcattgacaggtg-catgacagttcagctgc-----
shamisen shell     tcat-ggtttgacaggtg-cgtgacagtcagctccc-----
horseshoe worm     --at-gggttgacaggtg-cgtgacagttcggc--------
bootlace worm      --at-ggcttgacaggtg-cgtgacagttcagct-------
```

**Figure 23:** *Ssbp2* promoter. This matches two places in the scorpion genome and four places in horseshoe crab, all upstream of genes annotated as "Ssbp3-like".

```
human ZNF503           ----------------------------------------------------------------------------------------------------------------------ctcttttcccttcaagtcgca
chimaera znf503        -------------------------------------cgtccgccttttgttcgcatacaaattgagtttactctgagaacgatttatttccctatattgaatgcctcttttcaaa-agccgttctttttcaagtcaca
lancelet               aaaggggcacaaattgccgcctacttacag----tgtttcggcgttctttgttcgc-cgccaaacgggtttattt--ccaacgattta-tatcaccaattcaatgacgcttttcaaagcctttcttttttcaaatcaca
acorn worm             ---------------------------------------gactccctttgttgtcggacaaattgagctcacac--agaccgatttactttcggcaattcactggccttttcagaa------aactttttcaaattacc
sea urchin znf503      -------------------------------------cggtctctttgttgtcacacattaattgcgattatt---gggccgatttatctcggggaatcataccccctttcaaaag------ccgtttttcaaattact
sea urchin znf503-like -------------------------------------cggtctctttgttgtcacacattaattgcgattatt---gggccgatttatctcggggaatcataccccctttcaaaag------ccgtttttcaaattact
millipede              -------------------------------------------------------------------------------------------------------------------------------------------
scorpion               -------------------------------------------------------------------------------------------------------------------------------------------
scorpion               -------------------------------------------------------------------------------------------------------------------------------------------
scorpion               -------------------------------------------------------------------------------------------------------------------------------------------
tick                   -------------------------------------------------------------------------------------------------------------------------------------------
horseshoe crab         -----------------------------------------------------------------------------------------------------------------------------------------cg
horseshoe crab         -------------------------------------------------------------------------------------------------------------------------------------------
velvet worm            ---------------------------------------------------------cgcaaattgcgtttattt--tgaacgattag-tgtgaccaattcaatgtgtttt-----------attctttatttgctcgg
cactus worm            -------------------------------------------------------------------------------------------------------------------------------------------
abalone                -------------------------------------------------------------------------------------------------------------------------------------------
shamisen shell         agacggatacaaacttccatct-cctacaggatcggctttcaaattctttgttcacgcacaaattgcgtatactt--tcagcgattag-aattaccaattccgtgtggttt-----------atcctttatttgcctct

                                                       MEIS__________
                                           MafF____________          PKNOX__________           MEIS__________                  NFI_____________
human ZNF503           cccgc-cttg---taatcagcaacaaaa-gctgacataaatcacgggaggattgacaagagctgaca-aataactgattgattg-cggtcgaggaacattgacaattgatggaggggtggagatgcccaaagaaggggg
chimaera znf503        ccagg-cttg---taatcagcaacaaaa-gctgacataaatca-acgcagattgaca-gggctgaca-aataactgattgattg-cggtcggggaa-attgacagtctggggaggggaagg------aggggaggggag
lancelet               ccggaccatg---taatcaacaacaaaa-ggtgacatgaatca-gccaggattgaca-gccttgaca-ggagattgattgatgg-gggtctggat--atcgacggatggcgacatgccagg--tgattaa---------
acorn worm             ctagc-catg---taatcgcctacaaag-ggtgacataaatca-gacgcgattgaca-gccatcacg-ttggaatgattgatta-cgct--------------------------------------------------
sea urchin znf503      cgagc-catg---ttatcaacaacaaaacgctgacataaatca-gacggaattgaca-aggctgacaccgcatttgattgatcg-cag---------------------------------------------------
sea urchin znf503-like cgagc-catg---ttatcaacaacaaaacgctgacataaatca-gacggaattgaca-aggctgacaccgcatttgattgatcg-cag---------------------------------------------------
millipede              -----------------cgacaacaaaa-ggtgacataaatca-agtatgattgacgtgccgtgaca-ggcgcttgatggatga-ggtcct------------------------------------------------
scorpion               -------------------acaacaaaa-gctgacataaatca-accttgattggca-cttgtgaca-ggtgtgtgatagatgg-cgctgtaaac--catgacggctggcggcttgccaag--tcattag---------
scorpion               -------------------acaacaaaa-gctgacataaatca-accttgattggca-cttgtgaca-ggtgtgtgatagatgg-cgctgtaaac--catgacggctggcggcttgccaag--tcattag---------
scorpion               ----------------ttgacaacaaaa-gctgacataaatca-actgtgattggca-cttgtgaca-gccgcttgatggatcg-ccttgtagcc--tgcgacgcctggcatcttgccaag--tcatta----------
tick                   -------------------acaacaaaa-gctgacataaatca-tgcctgattggca-gctctgaca-acgggctgatggatag-cgttgtaaaa--gttgacagctggccccacgccaga--tcatta----------
horseshoe crab         tcgcactgta---ggcccgacaacacaa-cctgacataaatca-actgtgattgaca-cttctgaca-gatccttgatggatgg-tgttgcaaag--actgacaactggcggcttgccaaa--tcatta----------
horseshoe crab         ----------------tcaagcactaag-cctaacataaatca-gttcctactgaca-ctcttgact-ctcgactgatggatga-cgttgtaaac--cttgacagctggcactttgccaag------------------
velvet worm            ctatataatgtatacattaacaacaaaa-ggtgacataaatca-acgttgattgaca-gagctgaca-ggcgattgattgatcg-tagtctgaag--cgtgacggctggcggcttgccaag--ctatta----------
cactus worm            ---------------attactaacaaag-ggtgacataaatca-gttgtgattgaca-gccgacgca-ttgaagtgattgatgg-cg-----------------------------------------------------
abalone                ---------------attaacagcgcgc-ggtgacataaatca-agcttgattgacg-gacaggcca-tcaaaatgattgattgttggggcatga--ggtgacgtctggctagctgccaag--taatta----------
shamisen shell         cttca-agtgtatacacgaagaactaaa-gctgacataaatca-agcatgattgacg-tgcggctca-tcaacttgattgatca-cagtcggtgc--aa----------------------------------------

human ZNF503           gagggaggaaa
chimaera znf503        ggcagtggaaa
lancelet               -----aatgtc
acorn worm             -----------
sea urchin znf503      -----------
sea urchin znf503-like -----------
millipede              -----------
scorpion               -----tacatc
scorpion               -----tacatc
scorpion               -----------
tick                   -----------
horseshoe crab         -----------
horseshoe crab         -----------
velvet worm            -----------
cactus worm            -----------
abalone                -----------
shamisen shell         -----------
```

**Figure 24:** *Znf503* enhancer. This matches three places in the scorpion genome, two upstream of genes annotated as "Noc-like", and two places in horseshoe crab upstream of Noc-like annotations. (*noc* is an insect homolog of *Znf503/703*.)

```
                                                                      GC-box_
                                                              EGR_____________                                             DUX__________             ALX____
human ZNF503    gg---cctgacctgtcaggcggatta-----tcttcgagggagattaataggggaggcgggctgcaataataatcgttttgtttgatgtgacaactctgatagg-cgttgatttacttacaaactgataggcttttaatt
human ZNF703    tga-ccctgagccgtcaagaag-----------gaacagccagattaatggggtgggcgggctggggtaataatcg-tttgtttgatgtgacaagcctgatagg-cgttgatttacttacagactgatgggcttttaatt
chimaera znf503 ga---cctgacctgtcaagcggatta-----tcatacagagagattaatagggtgggcgggttgcaataataatcgttttgtttgatgtgacaagtctgatagg-cgttgatttacttacaaactgataggcttttaatt
chimaera znf703 tgaccccgagactgtcaagcga-----------agtcccggagattaatggggtgggcgggttaaagcaataatta-tttgtttgatgtgacaggcttgatggg-cattgatttagttgcggactgataggcttttaatt
lancelet        ta----taaacgtgtcagtaggattaccgaatgatggagagtgataaatggggtgggcggtctatgataataatcattttgtttgatctgacacttttgatagg-cgttgatttactttaaaactgataggcttttaatt
acorn worm      -------------------------------------------ataaatagtatgggcgtacatgggtaataatcttattgtccacgttgacggttttgattga-cgttagtttact---------acgtcgttttcata
velvet worm     ta----taaacgcgtcagccacttcactgattgatgcaagcagataaataggataggcggggcagccaaacaatcattctgtttgatttgacacgtctgatagaccgttaatttact-----------------------
shamisen shell  ------------------------------------caagtagattgatggagaaggcggggtgtctaaacagttcccttgtaccacttgacacgtctgatagg-gcttaatttacttaaagcct--ctgcgttccgatt

                ---
human ZNF503    gagcgcctccg-ccgagcccgagatgaaagggaagcggcg----------------------------------------gtg
human ZNF703    gagcacgccat-cctagtc--acttcaaag-----------------------------------------------------
chimaera znf503 gagcgcctacatccaagttttagatgaaaggaaagcggcg----------------------------------------gcg
chimaera znf703 gagaacctccaattccgaggcacatcaaag-----------------------------------------------------
lancelet        gagcgcctaag-gagcggc-gacttcaaacaagtcgaggacg--taagctgtctcttttggcggggcaacaaaacaatttccc
acorn worm      gcacaccgccg--agcggc-gacttcaaagaaatactccccactatagctgtct---------gagcaacaaaacaatttgcc
velvet worm     -----------------------------------------------------------------------------------
shamisen shell  actatcaaaacacggcggccaacatcaaaggcgtcctgcgtagcctaccgtctcgtttgggcaagggaacgtaacaatttgcc
```

**Figure 25:** *Znf503/703* enhancer

```
human      caggttggcagatctggagggctaatccaaacaa----ggcgcttgacttgtgaggtc-gcgagtctagacagagctggtggctcggcggctccttatctttccctggaatcagccgaagag
chimaera   caggttggtgtatctctgagcctaatcaaaacaa-------gtttgacttgtgaggtcctcacgtctagacagagctggactctcggcggctccttatctgagaacggaatcagcagagagg
acorn worm ----------------------aatcaaaacagttttgtttttttgacttgtgaggtctgtacgtctagacagaaagcgtcgcaaggcggcggcttatctcccttcggcatcaaactaa---
```

**Figure 26:** *Isl1* enhancer

```
human      gtctttaatgtaccattaactattgtctttacacaatatgggaactgt---aaagcatgacatgtgttataataaaacaca-ttttcaatg-atacacttggacttgagg-ctggggtgc-agcaaacaaaca
chimaera   gtctttaatgtaccattaactattgtctttgcacaatatgggaactgt---aaagcatgacatgtgttataataaaacaca-ttttcaatt-atacacttgtactggaag-ctctgatgcaagcaaacaaaca
acorn worm gcctttaatgttctctgggcctttgtacgttcacaatgtgggcacact---agagaatgacgtgtctcggataaaaacacacttttcaatt-atctagttatagtgtagg-ctgtgaagcaaacaaacaatca
sea urchin -------------------ccattgtcccttgacaatgtatgcacagctcgtgatgatgacgtctgccccataaaaacacacttttccattggtctaggtttcctgtaagcctttgaagtaaaca--------
```

**Figure 27:** *Ptf1a* enhancer

```
human SIX1    -agacttgtccccctggcatctcaggttaa--agaaggaggtccaggaaaatgt-ggtagagccattacaagaaacccgagccgcagtcatca--------
chimaera six1 tgaggtcggccctccagcttctcgggttaa--aggaggcgggcctgcaaaatgt-ggtagagccattacaagaaacccgagatccattcatcatgcactca
chimaera six2 --ggtcgaccccaccagcctctcgggttaggtagtgggagg-ttgtcaaaatacaagagcagccattagtagaaacgtgatatctgttcatca--------
lancelet      -agggtcggccctctgccggctcgggttgt--agggggagg-tttagaaaatat-ggtatagtcattaagcgaaatttgatacccattcatcatgcgc---
acorn worm    tcaggtcagccctctcgcgtctcgggttga--agtaggagg-tttaagaaatat-ggcatagtcattaagggtaattcgatacccattcacgatctagtca
```

**Figure 28:** *Six1/2* enhancer

```
human SP8    gaggccgcagattggctggcc--gcgcctccgcggcacttcaaagccggagagggagctacaattg--tggtggaat-gggcggaactgatcatt-gtattgcacacct
chimaera sp8 -----ccctgattggctgccg--gggtctgagcggtacttcaaaggcgagagaggcgctacaattg--cggttgaatggggcggaactgatcatt-gtgttgcacacct
chimaera sp9 -------ctgattggccgagggctcggcacagcggcacttcaaagcccaca--cccgctacaatta--gggtcgagt------------------------------cg
lancelet     gagctcgctgattggtcgagag-gggcctgagcgggccttcaaagggaagc--cacaccacaattatatggtcaaat-gggcggaaccgaccatttgtattgtacacct
acorn worm   --------tcattggtggatca-gccgataggcggctcttcaaagcccttt-gcaagcggcaattatcgggccaaat-gggcggacctggtcatttgcat---------
```

**Figure 29:** *Sp8/9* promoter

```
human ZFP36L1    gcttgt---------------------------cactgcacat-caatataaaaaa---gcttatttaacttatcaaaacgtattt-attgccaaactatgcttt-----tttttgttaattttgt------tcatattt
human ZFP36L2    tttttagctgttttgagttgattcgcaccactgcaccacaact-caatatgaa------aactatttaactta------tttatta---tcttgtgaaaagtata------caatgaaaattttgt-tcatactgtattt
chimaera zfp36l1 gcttct---------------------------cactgcacacacaacacaaaaaaattgcttatttaacttatc-gaacatattt-attgccaaac--------------atatgcattttttgt------ttgtattt
chimaera zfp36l2 tttttagctgttgagtttttgatggcccactggtggcccagct-cgctataa-------aactatttaactta------tttatta---tctcg-agaaagtata------cattggtaa-tttgt-tcatactgtattt
lancelet         tattta-----------------------------------------------------acttatgtgactta------tttatttgaaccttatacgtaacgttatgcatttttggtaaatttattatgctcgttatgt
acorn worm       --------------------------------------------------------------tatttaagttaacaggccttattt---atttgaagtaagcatt------ttttggtaaatttataatgcctcctattt

human ZFP36L1    atcgggatgacaaat---ccataga-atatatt---cttttatg-ttaaattatgatcttca-tattaatcttaaaattttgtgacgtgtctttttcct-----------------------------------------
human ZFP36L2    atcaagtatgatgaaaagcaatagatatatattcttttattatg-ttaaattatgattgccattattaatcggcaaaatgtggagtgtatgttcttttcacagtaatatatgcc--------------------ttttgt
chimaera zfp36l1 atcaaaaatgatgaa---ccataga-gtatatt-ttcttttatgtttaaattatgatcttc----ttaatcttaagattatgaggtcctttttttggtc-----------------------------------------
chimaera zfp36l2 atcgcgtatgatgaaaa-caatagatatatattcttttattatgtttaaattatgattgccattattaatctgcagaatgtggagtgtatgttgttttcacagtaatatatgccttttttgttgggtttttttgttttgt
lancelet         acgaga--cgatgagag-tgatagatatttatt---atattatgtttaatttatgattgtt--tgtgaatcacaggattggagtgtgtgtactatgtgtatggtaatatgtg----------------------------
acorn worm       atcgggttccatgtggg-acatagatatatatt-ataaactttg-ttaaattatgattgtgaattgaaatggacagagactggagtgt----------------------------------------------------

human ZFP36L1    ----------------tttttt
human ZFP36L2    aacttcac-ttggttattttat
chimaera zfp36l1 ----------------tttttt
chimaera zfp36l2 aacttcactttggttattttat
lancelet         ----------------------
acorn worm       ----------------------
```

**Figure 30:** *Zfp36l* 3′-UTR

```
human EBF1     gggcccggtatggatgattaggtt------tcgggctttgaattgttgacaaggcaggggtaatgaattgcgggacactggctcctgcatttccccctcattagtcagtattagctacattaatagtaataatt------
human EBF3     -------------------------------------------------------------------------------------cacatgtgcccctcattaagctgcattacctgcattaaaactaataattgccact
chimaera ebf1  ggcccagccgaggatgattagctttgtggatttgactgtgatttgttgacaaggcagagg-gatgaattgtgggac--ttgagcaagtgtgtgactctcattattcagtattagccgcattaatagtaataatt------
chimaera ebf3  ------------------------------------------------------------------------------ttgcacacatgtgcccctctcattaagctgcattacctgcattaaaactaataattgccact
lancelet       ----------------------------------------------------------------------------------ttgcaggcatgcctctcattaagttacatcacatgcattagaagtaatcattcctgct

human EBF1     ---------ggatatattcatcatttaaaatgttttaataaatgtttggacagtacccgatcagagctgtcaaa-------
human EBF3     cagagctgtgcgcccattaatcatttataatgttttaataagtttttaatcaggacctaa--aaggctagagagccgcata
chimaera ebf1  ---------acatgaattaatcatttataatgttttaataaatgtttggtcagcacctgatctacactgtcaaa-------
chimaera ebf3  cagctctgtgctcacattaatcatttataatgttttaataagtttttaatcaggatttga--aaggctagaaagct-----
lancelet       cacctccaagcccccattaatgatttaccaggtcctaatcaggttttgatgggagtgtac--ggggctagcgagctgtgca
```

**Figure 31:** *Coe* intron element 2

```
human IRX1     ---ttttctacttgaacatgaaataatgattttttgctcatataattacaaagctaacatctgatagagtcagcatccaggggggattatcgcatgcatgagtattagacctcattaccagcttgactgcaaaattatta
human IRX3     tcattttctatttgaacatgaaataatgattttctgcccgtctaattacaaagccaacatctgatcaagccagcatccagaggggattatcgcatgcacgagtattagacctcattaccagcttgagtgcaaaattatta
human IRX4     tcatttttctgctgaacatgaaataatgattttctgccagtataattacaaagctaacatctgatagagtcagcatccaggggggattatcacaggcatgagtattagacctcattaccagcttgactgcaaaattatta
human IRX6     ----ttttttcctgaacatgaaataatgattttctgcccgtctaattacaaagccaacatctgatcaagccagcatccagaggggattatcgcatgcacgagtattagacctcattaccagcttgagtgcaaaattatta
chimaera irx1  ----tttctacttgaacatgaaataatgattttttgctcatataattacaaagctaacatctgatagagtcagcatccaggggggattatcacatgcatgagtattagacctcattaccagctcgactgcaaaattatta
chimaera irx3  tcattttctatttgaacatgaaataatgattttctgccagtctaattacaaagccaacatctgatcaagccggcatccagaggggattatcacatgcacgagtattagacctcattaccagcttgagtgcaaaattatta
chimaera irx4  ---tttttttgctgaacatgaaataatgattttctgccagtataattacaaagctaacatctgatagagtcagcatccaggggggattatcacatgcatgagtattagacctcattaccagcttgactgcaaaattatta
chimaera irx6  ----ttttttcctgaacatgaaataatgattttctgccagtctaattacaaagccaacatctgatcaagccggcatccagaggggattatcacatgcacgagtattagacctcattaccagcttgagtgcaaaattatta
lancelet IrxA  ----------------------------------------------------------------------------------------------------agaattataactaattataagcatgactgccacaatatta
lancelet IrxC  ----------------------------------------------------------------------------------------------------agaattataactaattgtaagcatgactgccacaatatta

human IRX1     catcttgtaattggatggtgag--------atttatatacagattgg----ccgggtt-tctgtaagattgtaattacaagcttcattggtttgctacttatcgga-acttcacagaga----aaacaactggtaaaatg
human IRX3     catcttgtaattggatggtgag--------atttatatacagatcgg----cccggtt-tctgtaagattgtaattacaagcttcgttggttagctacttatcagaactttgcaggaaaaca-aaacaaatgactgagga
human IRX4     catcttgtaattggagggtgag--------atttatatacagattgg----ccgggtt-tctgtaagattgtaattacaagcttcgttggtttgctacttatcaca-cggtgcaggaggcggc--aaaaacaagc-agag
human IRX6     catcttgtaattggatggtgag--------atttatatacagatcgg----cccggtt-tctgtaagattgtaattacaagcttcgttggttagctacttatcact-cagcacagaagag---aggaaaataagc-aaca
chimaera irx1  catcttgtaattggatggcgag--------atttatatacagattgg----ccaggtt-tctgtaagattgtaattacaagcttcattggtttgctacttatcgga-acttcacagaga----aaacaactggtaaaatg
chimaera irx3  catcttgtaattggatggtgag--------atttatatacagatcgg----cctggtt-tctgtaagattgtaattacaagcttcgttggttagctacttatcagaactttgcaggaaaaca-aaacaaatgactgagca
chimaera irx4  catcttgtaattggatggcgag--------atttatatacagattgg----cccggttctctgtaagattgtaattacaagcttcgttggtttgctacttatcaca-cagtgcaggaggaggcaaaaaaacaagc-agag
chimaera irx6  catcttgtaattggatggtgag--------atttatatacagatcgg----cctggtt-tctgtaagattgtaattacaagcttcgttggttagctacttatcaca-cagcacaggaga----aaaaaaataagc-aaca
lancelet IrxA  caactgataattggaaggttggatggcttcatttgcatgtagagctgagacctgggtt-cccaccaga---caattatagctttggttagagggagcctcttggct-ccatgcaggatgctgctaca---------cacc
lancelet IrxC  catctgataattggaaggttggatggcttcatttgcatgtagagctgagacctgggtt-cccaccaga---caattatagctttggttagagggagcctcttggct-caatgcaggatgctgctaca---------cacc

human IRX1     aaatgagcatttcattaacctattgccttatcatgggaaaacctgtgtaacagccgtgtatgcattaacataaatag---------
human IRX3     aaacgagcatttcattaacctgtaacccgagcgcgaggg-----------------------------------------------
human IRX4     ggaaggccatttcattaagctgtaaccctatcaagggaggagcgctgtaacgtccctgtatttattaggccaaacag---------
human IRX6     gaaaaggcatttcattaacccataaccctatcaagggaaagccactgtaacacccatgtctgcattagggagaacagctaaattca
chimaera irx1  aaatgagcatttcattaacctatcgccttatcatgggaaaacctgtgtaacgttcatgtatgcattaacataaatag---------
chimaera irx3  aaatgagcatttcattaacctgtaaccctatcatgggggaacctgtgtaacattcatgtatgcattaaggaaaatgggttcatcca
chimaera irx4  gaaagaccatttcattaacctgtaaccctatcaagggaggagcactgtaacatccctgtatttattaggccaaacag---------
chimaera irx6  gaaagagcatttcattaacccgtaaccctatcaagggagaactgctgtaacatccatgtttgcattagggagaacag---------
lancelet IrxA  acaagaccactttattaacc------------------------------------------------------------------
lancelet IrxC  acaagaccactttattaacc------------------------------------------------------------------
```

**Figure 32:** *Irx* enhancer

```
human NR2F1    ----------------------aaacaagaattaggggaaaataacattttccaaataattataaaaaattgtcctgtgtctatgtatctatatctg---ttttgta-tttttttctggttccaaaccagatt-tcctgt
human NR2F2    -----------------------------------------------ttttccaaattatt------aattgtcctgtgtctatgtacctctagctgttcttttttg-tacttttctggttccaaaccagtttattctgt
chimaera nr2f1 gcgcatttaaaagacaga---atcaagagattagggggaaaataacattttccaaataattat---aaattgtcctgtgtctatgtatctatatctg---ttttgtattttttttctggttcaaaaccaggtt-tcgtgt
chimaera nr2f2 gtgcatttaaacaagaatttcaaaattaggaggaaagaaaagtaacattttccaaattattat---aaattgtcctgtgtctatgtatctatagctg---ttttgta-ttttttcctggttccaaaccagtctcctctgt
lancelet       ---------------------------------------------------------------------------tgtgtctatgtatctt--gctg---ttttata-----tacatggttccaaaccaaacg-ttaagt

human NR2F1    gatt-ctatactaataattttttgatataaccctttgcttcttataatgagtgcgatatatgttgtcgaggc--tgttcttcaagaattaaaattgaagtga----aaatttaaacaaaaa
human NR2F2    ggtt-ctataata--ag-ttttgatataatc-ttggcttctt-----aaaaactgtgtatcattaaaatat-atgttctgcaagaattaaaactgagtccatgaaaataccataggaaga
chimaera nr2f1 gattcctatactaataattttttgatataaccttttgcttcttataatgagtgagatttatgtcgtcaaagctatgttctccaagaattaaaattgaaatca-------------------
chimaera nr2f2 gatt-ctctactaataa-ttttgatataacc-attgcttcttat--cagatacggtgtatcacgtaaat---atgttctgcaagaattaaaactgaattcactaaaatgttagataaaaa
lancelet       gatt-tactaataataa-ttttgatataacc-----------------------------------------------------------------------------------------
```

**Figure 33:** *Nr2f* 3′-UTR
